# Modulation of blood-tumor barrier transcriptional programs improves intra-tumoral drug delivery and potentiates chemotherapy in GBM

**DOI:** 10.1101/2024.08.26.609797

**Authors:** Jorge L. Jimenez-Macias, Philippa Vaughn-Beaucaire, Ayush Bharati, Zheyun Xu, Megan Forrest, Jason Hong, Michael Sun, Andrea Schmidt, Jasmine Clark, William Hawkins, Noe Mercado, Jacqueline Real, Kelsey Huntington, Mykola Zdioruk, Michal O. Nowicki, Choi-Fong Cho, Bin Wu, Weiyi Li, Theresa Logan, Katherine E. Manz, Kurt D. Pennell, Bogdan I. Fedeles, Alexander S. Brodsky, Sean E. Lawler

## Abstract

Glioblastoma (GBM) is the most common malignant primary brain tumor. GBM has an extremely poor prognosis and new treatments are badly needed. Efficient drug delivery to GBM is a major obstacle as the blood-brain barrier (BBB) prevents passage of the majority of cancer drugs into the brain. It is also recognized that the blood-brain tumor barrier (BTB) in the growing tumor represents a challenge. The BTB is heterogeneous and poorly characterized, but similar to the BBB it can prevent therapeutics from reaching effective intra-tumoral doses, dramatically hindering their potential. Here, we identified a 12-gene signature associated with the BTB, with functions related to vasculature development, morphogenesis and cell migration. We identified CDH5 as a core molecule in this set and confirmed its over-expression in GBM vasculature using spatial transcriptomics of GBM patient specimens. We found that the indirubin-derivative, 6-bromoindirubin acetoxime (BIA), could downregulate CDH5 and other BTB signature genes, causing endothelial barrier disruption in endothelial monolayers and BBB 3D spheroids *in vitro*. Treatment of tumor-bearing mice with BIA enabled increased intra-tumoral accumulation of the BBB non-penetrant chemotherapeutic drug cisplatin and potentiated cisplatin-mediated DNA damage by targeting DNA repair pathways. Finally, using an injectable BIA nanoparticle formulation, PPRX-1701, we significantly improved the efficacy of cisplatin in patient-derived GBM xenografts and prolonged their survival. Overall, our work reveals potential targets at the BTB for improved chemotherapy delivery and the bifunctional properties of BIA as a BTB modulator and potentiator of chemotherapy, supporting its further development.

## Introduction

The effective treatment of brain malignancies such as glioblastoma (GBM), remains a critical challenge in the neuro-oncology field. GBM is the most common malignant primary brain tumor, representing ∼15% of all central nervous system (CNS) neoplasms ^1^. Median survival is 15-18 months and less than 10% of patients survive beyond 5 years after diagnosis ^2^. The standard of care involves maximal safe surgical resection, followed by radiotherapy and alkylating chemotherapy with temozolomide (TMZ). This inevitably leads to the development of untreatable recurrent disease. The major challenges in GBM therapy are 1) its invasiveness, which prevents complete surgical resection, 2) high levels of intra-tumoral molecular and cellular heterogeneity, 3) a cancer-promoting tumor microenvironment (TME), and 4) the presence of the blood-brain barrier (BBB) and blood-brain tumor barrier (BTB), that limit drug entry.

The BBB maintains homeostasis of the central nervous system (CNS) for its proper functioning ^3^. In an oncogenic context, the BBB responds to cues by the cancer cells, which promote the formation of new blood vessels and the BTB. The BTB is a distinct and heterogeneous biological entity, resulting from cellular interactions between brain-tumor cells newly formed blood vessels, and the pre-existing BBB ^4^. Molecular characteristics that define the impermeability of the BBB, such as tight junction and adherens junction formation, high eflux pump expression and non-fenestrated endothelium, are compromised in brain tumors mainly due to hypoxic/angiogenic conditions, which also promote tumor growth, migration and invasion ^5^. Regardless of the disruption of these brain-protecting BBB properties, non-BBB penetrant drugs still do not penetrate GBM tissue efficiently. This is supported by studies suggesting that the BTB is highly heterogeneous ^6^, with some regions maintaining “healthy” BBB features that protect GBM cells from anti-neoplastic agent accumulation.

Many biological features of the BTB are poorly understood, especially its molecular and cellular composition, and identification of target molecular pathways that could render the BTB permissive to chemotherapy uptake. Strategies to improve drug delivery to GBM include focused ultrasound (FUS) ^7,8^, convection-enhanced mediated delivery (CED) ^9^, optogenetics ^10^, systemic administration of drug-loaded nanoparticles ^11^ and drug-conjugated cell-penetrating peptides ^12^, with most of these options showing promising pre-clinical results. These strategies rely on physically overcoming the BBB/BTB, and have advantages of controlled release, preservation of drug stability and drug delivery at selected anatomical sites. In addition to these approaches, the identification of compounds that could target molecular elements that selectively regulate BTB permeability, but not healthy BBB, would enable mechanistic control over the biological processes involving the BTB/GBM tumor interactions, and could be used to potentiate intra-tumoral drug penetration. Moreover, if these compounds could simultaneously hinder tumor development and synergize with chemotherapeutic regimens, this would be potentially useful in the clinic.

Previously, we identified the anti-invasive and immunomodulatory properties of the indirubin-derivative 6-bromoindirubin acetoxime (BIA) in GBM and showed some benefit in murine GBM models ^13^. We also developed a BIA-loaded nanoparticle formulation, PPRX-1701, which was well-tolerated, and able to reach intracranial brain tumors in mouse models ^17^. Indirubins are bisindole alkaloid compounds used as a component of traditional Chinese medicine for the treatment of proliferative disorders and auto-immune conditions. Indirubin is a component from the *Indigo naturalis* extract ^14^. BIA is widely known as a GSK-3 inhibitor ^15^, but several other kinases have been found to be inhibited by this compound, including cyclin-dependent kinases and Src-family kinases ^16^.

Herein, we report that BIA has significant effects on BTB permeability by reducing the expression of BTB signature genes, including the tight junction protein CDH5 (VE-cadherin). BIA treatment increased cisplatin accumulation in tumor tissue in mouse tumor models, but not in healthy brain, and enhanced the cytotoxic capacity of cisplatin. BIA in combination with cisplatin prolonged survival of xenograft GBM models. Together, our work provides evidence of potential candidate targets at the BTB and the use of BIA for improved drug delivery and chemotherapy potentiation in GBM.

## Results

### Identification of GBM tumor endothelium-associated genes (BTB-genes) via *in silico* screening

The BTB represents an obstacle to therapeutic drug delivery and remains a poorly defined component of GBM biology. Thus, to identify molecular signatures of the GBM vasculature for targeting of the BTB, we performed an *in silico*-based approach by accessing bulk RNA-sequencing data from The Cancer Genome Atlas via the cBIO portal (**Figure 1A**). We used Spearman’s rank correlation to select GBM genes that showed co-expression with four well-characterized endothelial cell markers defined by Dussart et al (2019) for screening brain tumor-associated vasculature ^17^: *CD31*, *CD34*, *VWF* and *CLEC14A*. (**Supplementary Table 1**). With this approach, we identified a signature comprising 12 GBM tumor endothelium-associated genes (BTB-genes) with increased expression in GBM tumors above non-tumoral tissue (**Figure 1B**). This BTB signature included angiogenesis-associated genes such as *ACVRL1 (ALK1)*, *CD93, ENG* and *PDGFRB*, as well as endothelial cell-adhesion endothelial genes *PCDH12*, *ROBO4*, *ESAM* and *CDH5*. These genes present abundant expression at microvascular proliferation regions in the tumor (**Figure 1C**). Gene Ontology analysis revealed their primary involvement in vasculature system development, morphogenesis and cell migration (**Figure 1D**). STRING network analysis showed that the genes form significant interaction within the network (PPI enrichment p-value: <1.0e-16). With the exception of *MYO1B* all genes were interconnected (**Figure 1E**), suggesting strong functional relationships within the GBM BTB signature gene set.

**Figure. 1.**
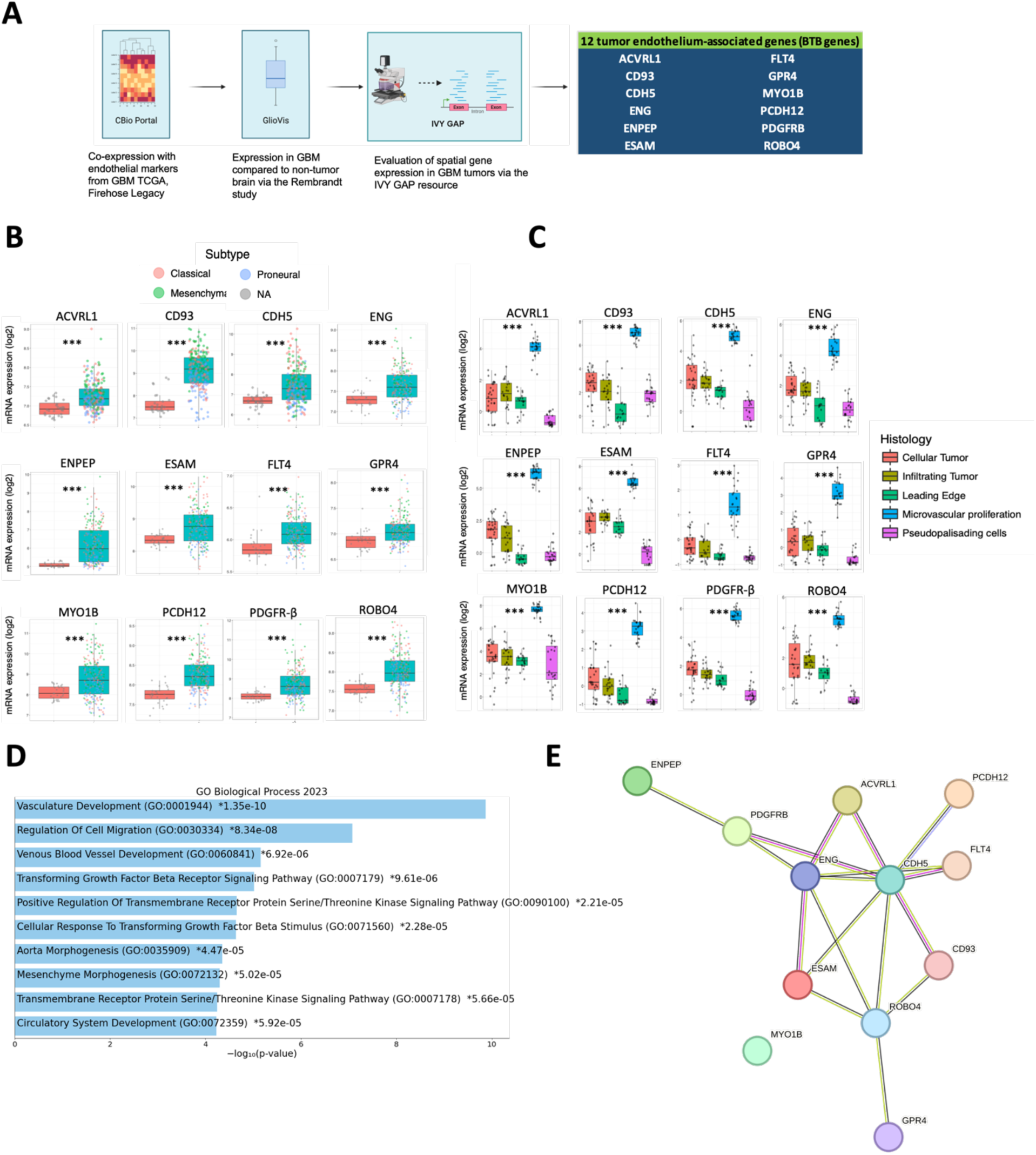
Identification of tumor-endothelium associated (BTB-genes) using bulk and spatial RNA-sequencing datasets from GBM clinical samples. (**A**) Workflow of identification and filtering of genes associated with tumoral vasculature in GBM. Using a gene expression correlation tool, cBIO, top-50 genes that co-expressed with CD31, VWF, CD34 and CLEC14A were selected, and their expression in tumor above normal brain examined in the Rembrandt dataset using the GlioVis visualization tool, and evaluated regional expression using the IVY GAP atlas resource. (**B**) Gene expression of 12 tumor endothelium-associated genes (BTB-genes) identified in the cBio Portal following the workflow shown in (**A**), *** p-value<0.001 by Tukey’s Honest Significant Difference test. Individual values are colored by GBM subtype Classical, Mesenchymal or Proneural, NA indicates unknown sample information. (**C**) Regional expression of the 12 BTB-genes in GBM using the IVY GAP resource, *** p-value<0.001 by Tukey’s Honest Significant Difference test. (**D**) Gene ontology analysis (GO Biological Process 2023) of the 12 BTB-genes using EnrichR software. Biological processes are ranked by p-values, which are indicated next to the GO designation. (**E**) STRING network analysis on the 13 identified BTB-genes. 12 nodes, 17 edges, average node degree 2.83. PPI enrichment p-value 1e^-16.

### Spatial transcriptomics of GBM patient samples confirm *CDH5* upregulation in tumor associated endothelium

The STRING analysis showed that *CDH5* (VE-cadherin, CD144) may represent a central hub in the BTB signature gene network. CDH5 is a calcium-dependent adherens junction protein with a fundamental role in maintaining BBB integrity. To further investigate its potential role in the BTB of GBM, we re-analyzed spatial transcriptomics data from malignant glioma tissue samples from Ravi et al (2022) ^18^ and confirmed the expression of CDH5 in comparison to matching non-tumorigenic brain cortex of samples UKF_242, UKF_248 and UKF_334 (**Figure 2A**). This data showed considerable spatially distributed expression of CDH5 above cortex controls. CDH5 is clustered across the tumor (**Figure 2B**), and highly expressed in clusters enriched in vascular markers such as PECAM1 and VWF (**Figure 2C**). CDH5 was enriched in Biological Process GOs related to (**Figure 2D**) Vasculature Development and Vascular Process of the Circulatory System (**Figure 2E**), supporting its involvement in vascular processes in GBM. Moreover, we identified a list of 61 additional genes regionally co-expressed with CDH5 (**Supplementary Table 2**), which includes genes such as *CCL2* ^4^ and *WNT7B* ^19,20^, which have reported roles in the BTB. This data supports the notion that CDH5 is highly expressed in tumoral vasculature and may be relevant to modulate BTB properties.

**Figure 2.**
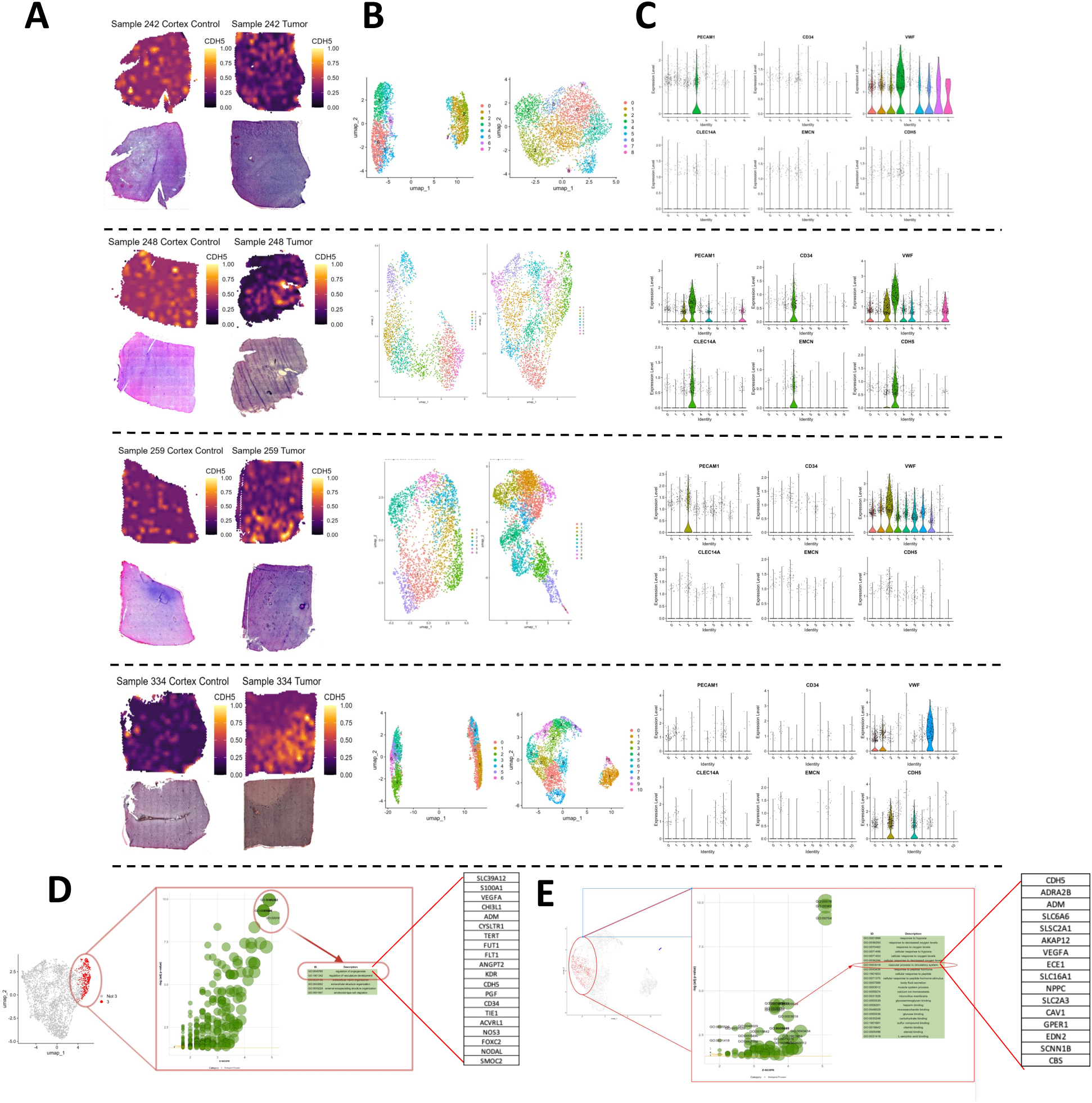
Spatial transcriptomic analysis of GBM patient samples confirms *CDH5* upregulation in tumor associated endothelium. (**A**) Surface plots of CDH5 expression from spatial transcriptomic performed on GBM tumors and non-tumorigenic cortex from cortex and tumor tissues. (**B**) Spatial UMAP plot of CDH5 expression in cortex (left) and tumor (right) showing specific clusters per sample. (**C**) Violin plots indicating endothelial cell marker expression and CDH5 from tumor tissues and clustered according to spatial clustering from (**B**). (**D**) Gene Ontology (GO) indicating associated pathways with CDH5 gene expression in cluster 3 of sample UKF_248 and (**F**) clusters 2 and 5 of UKF_334. GO biological processes highlighted in red indicate vasculature development and vascular processes of the circulatory system as pathways enriched for *CDH5*. Top 20 gene lists for highlighted GO pathways are shown. Data was obtained and re-analyzed from Ravi *et a*l., (2022) using the SPATA2 package from R-studio.

### BIA targets angiogenesis and BTB-related transcriptional programs in brain endothelial cells

Previously we demonstrated BIA has anti-angiogenic effects in murine intracranial models of GBM^13^. This led us to investigate the transcriptional alterations associated with BIA treatment of brain endothelium. Bulk RNA-sequencing analysis of a well-characterized human brain microvascular endothelial cell line, HCMEC/D3, treated with BIA showed considerable transcriptional dysregulation. The top 15 differentially expressed genes (DEGs) are displayed according to significance in a heatmap (**Figure 3A**). Interestingly, CDH5 was one of the most downregulated genes upon BIA treatment (−3.06-fold, log2). Gene Ontology (GO) analysis of significantly upregulated (862 genes) and downregulated (652 genes) differentially expressed transcripts revealed that BIA mostly induced expression of genes in processes related to amino acid transport. BIA also decreased expression of genes involved in annotated processes of cell migration, motility, angiogenesis, and endothelial proliferation, as well as nitric oxide synthesis and pathways of receptor tyrosine kinases (**Figure 3B**). Volcano plot analysis (**Figure 3C**) of log2 fold-change vs. p-value significance of downregulated DEGs highlights *CDH5* and other angiogenesis-related genes such as *MMRN2*, a direct interactor with CDH5, CD93, ACVRL1, KDR, SMAD6 and S1PR3. BIA also promoted expression of genes such as *PHGDH*, *AXIN2*, *TCF7*, *VLDLR* and *VEGFA*. Showing that BIA has broad effects on genes involved in diverse pathways. We then examined dysregulated genes that are potentially involved in BTB biology by focusing on BBB permeability/integrity and in biological functions of angiogenesis (**Figure 3D**). Indeed, BIA modulates 8 of our 12 BTB signature genes we identified in the *in silico* screening from clinical samples (**Figure 3D**, highlighted). Most of these genes were downregulated by BIA, except *PCDH12*, which increased its expression. This finding suggests that BIA targets the expression of BTB-associated transcriptional programs in brain endothelial cells. Our spatial transcriptomic analysis of sample UKF_248 showed the increased expression of these BTB-genes in GBM clinical samples above cortex controls (**Figure 3E**) and spatially co-expressed by clustering analysis (**Figure 3F**). These genes were expressed across tumor samples (**Supplementary Figure S1**) and indicate the relevance of these pathways in GBM.

**Figure 3.**
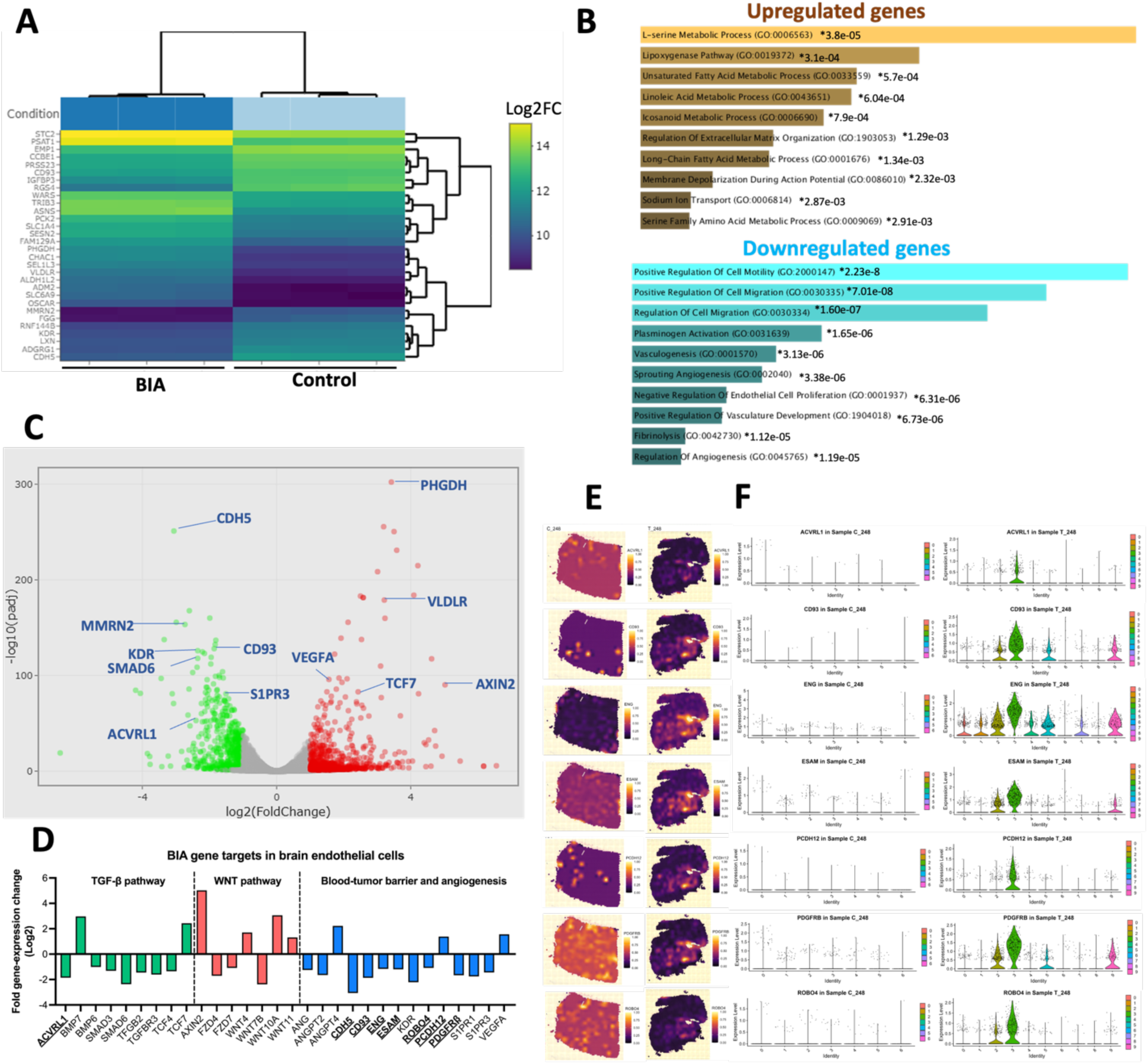
BIA modulates BTB-associated genes and cell motility, vascular development, angiogenesis and L-serine metabolism in brain endothelial cells. **(A)** Heat-map generated from the top-15 upregulated and downregulated genes (Log2FC) from Bulk-RNA sequencing analysis performed on HCMEC/D3 cells treated with BIA (1 µM, 24hrs). **(B)** Gene ontology analysis of >1.5 (Log2FC) significantly upregulated and downregulated genes by BIA in brain endothelial cells from **(A)**. Biological processes are ranked by p-values, which are shown next to the GO designation. Analysis performed using the EnrichR software. **(C)** Volcano plot analysis from all the upregulated and downregulated genes by BIA. Labels on genes related to angiogenesis, TGF-β and WNT pathways are highlighted. **(D)** Gene expression fold-change (Log2) levels of dysregulated genes by BIA related to the TGF-β and WNT pathways, angiogenesis and the tumor vascular associated genes (BTB-genes, highlighted). **(E)** Spatial expression of 7 of the 12 BTB-genes regulated by BIA *vitro* highlighted in **(D)** for non-tumor (left) and tumor (right) tissues from sample UKF_248. **(F)** Clustered gene-expression of UKF_248 showing the BTB-genes from (E) in non-tumor (left) and tumor tissue (right).

### BIA disrupts barrier formation and increases permeability in BBB models *in vitro*

Given the prominence of CDH5 in the BTB transcriptome, and its known role of maintaining vascular barrier integrity, we focused our efforts in further characterizing CDH5 expression in the BTB upon BIA treatment. Immunofluorescence (IF) staining of CDH5 showed a marked decrease at the membrane periphery in endothelial cells in vitro after treatment with BIA (**Figure 4A** and **Supplementary Figure 3A**). We observed a marked reduction in the BBB tight junction molecule ZO-1, but no difference in levels of Claudin-5. BIA decreased levels of CDH5 mRNA in brain endothelial cell lines, which declined for up to 48 hours following BIA treatment (**Supplementary Figure 3B**). Interestingly, BIA also reduced the expression of CDH5 in G34 GBM cells, with simultaneous decline of WNT7B and S1PR3 expression, suggesting that BIA can modulate these endothelial barrier-related molecules in the tumoral context as well and is not restricted to vascular cells only (**Supplementary Figure 3C**). Protein levels of CDH5 reached maximum reduction at 12 hours post-BIA treatment, and remained downregulated for a further 48 hours (**Supplementary Figure 3D**).

**Figure 4.**
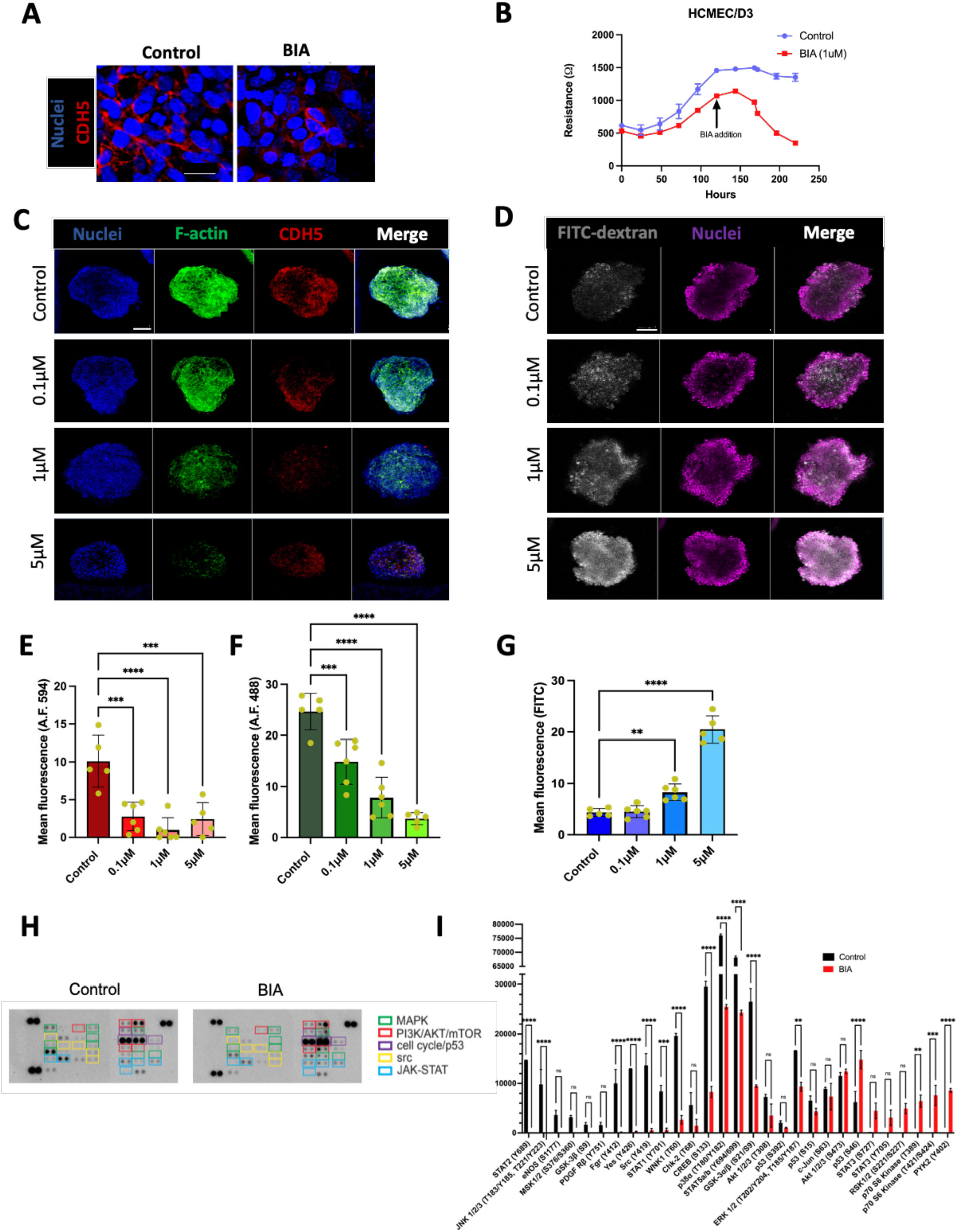
BIA prevents barrier formation by brain endothelial cells *in vitro* and increases dextran uptake in a three-dimensional BBB spheroid model. (**A**) Immunofluorescence staining of CDH5 (red) and nuclei (blue) in HCMEC/D3 cells treated with BIA (1 μM, 24hrs). Scale bar=20 μm. (**B**) TEER values of HCMEC/D3 treated with BIA upon monolayer confluence. Time-point of BIA addition is indicated. (**C**) Immunofluorescence images of BBB spheroids treated with indicated doses of BIA for 72 hours. Staining of F-actin (green), CDH5 (red) and nuclei (blue). Maximal projection intensity is shown from z-stack images (50 μm depth, 20 layers). Scale bar=100 μm. (**D**) FITC-conjugated dextran (70kDa) permeability assay in BBB spheroids. Dextran (gray) and nuclei (purple) are shown. Scale bar=100 μm. Mean fluorescence quantification of (**E**) CDH5, (**F**) Phalloidin and (**G**) FITC-dextran from images in (**C**) and (**D**) using the ImageJ software. Data shows mean and standard deviation, n=4-5. Ordinary one-way ANOVA test. ** p=0.0028, *** p=0.008, **** p<0.0001. (**H**) Human phospho-kinase array of HCMEC/D3 cells exposed to 1 μM BIA for 24 hours. Highlighted wells related to indicated pathways. Samples were analyzed in duplicates. (**I**) Quantification of signal by ImageJ of dot-blot shown in (**H**). Mean and standard deviation of duplicates are shown. Two-way ANOVA analysis was performed. ** p=0.0015, *** p=0.005, **** p<0.0001.

To understand whether BIA might alter barrier formation properties in brain endothelial cells, we performed trans-endothelial electrical resistance (TEER) analysis of monolayers of HCMEC/D3 cells. Treatment with BIA led to a marked decrease of barrier integrity (**Figure 4B**). Moreover, addition of BIA 24 hours after plating endothelial cells completely prevented barrier establishment (**Supplementary Figure 3E**). These effects occurred from 100 nM to 10 μM BIA (**Supplementary Figure 3F**), confirming that BIA can disrupt BBB integrity *in vitro*.

We next tested the effects of BIA on vascular permeability measuring dextran uptake using an in vitro multicellular BBB spheroid model ^21^. In this experiment BIA decreased the expression of CDH5 in a dose-dependent manner (**Figure 4C and 4E**) as shown by IF staining. F-actin was also reduced considerably (**Figure 4C** and **4F**). Incubation of a fluorescent-dextran (70kDa) with the BBB spheroids treated with BIA showed a dose-dependent increase in permeability (**Figure 4D and 4G**).

To understand whether the effects we observed are a consequence of endothelial cell death, we screened for apoptosis via flow cytometry, which did not show late apoptosis/necrosis at any of the BIA concentrations used in comparison to a cisplatin control (**Supplementary Figure 4A**). Cellular ATP-content was reduced up to 30% in the ∼1-5 μM BIA range, and ∼50% and above for HBMEC cells treated at the same concentrations (**Supplementary Figure 4B**), indicating that BIA impacts endothelial cell metabolism. Visual assessment of HCMEC/D3 cells treated with BIA did not reveal signs of apoptosis or necrosis, but an elongated phenotype with long filipodia (**Supplementary Figure 4C**). Cell cycle analysis via flow cytometry showed a slight decrease in proportions of cells in G1 and G2/M phases, indicating that BIA affects endothelial cell proliferation but does not induce cell death at the concentrations tested (**Supplementary Figure 4D**). Indeed, cell counts of endothelial cells treated continuously with BIA showed cell numbers decreased significantly after 4 days post-treatment (**Supplementary Figure 3E**).

### BIA targets several kinases in brain endothelium *in vitro*

BIA is a broadly selective protein kinase inhibitor ^15,22^. To elucidate the kinase signaling pathways altered by BIA that could be involved in barrier modulation, we treated HCMEC/D3 cells with BIA and performed phospho-kinase array profiling (**Figure 4A and 4B**). We observed a decrease of activating phosphorylation in members of the MAPK family (p38α, JNK1, MSK1/2 and ERK1/2), SRC family (SRC, YES, FGR) and transcription factors at activator sites (CREB, STAT1, STAT2, STAT5a/b and c-JUN). The MAPK and SRC pathways are known to control endothelial transcriptional programs through CREB and other transcriptional regulators ^23–25^. On the other hand, we observed increased phosphorylation of STAT3 at S727 and Y705, and in p70 S6 kinase, which suggests activation of the mTOR pathway. Finally, secretome analysis of HCMEC/D3 cells treated with BIA indicates a pro-inflammatory secretion profile with an increase of cytokines such as TNF-α, IFNγ, IL-17A, IL6, IL-1β, prolactin, CCL8 and CCL4, among others (**Supplementary Figure 5A**). Whereas significant downregulation was seen to occur for CCL2 (**Supplementary Figure 5B**). Overall, our results indicate that BIA operates at different cellular signaling levels that induce diverse biological changes in brain endothelium, which might be required to induce the endothelial barrier disruption phenotype observed.

### BIA increases intra-tumoral drug accumulation in murine intra-cranial models of GBM

To understand whether BIA could also increase permeability in the BTB in the context of GBM *in vivo,* we implanted patient-derived GBM cells (G30) in nude mice and treated them with BIA and administered sodium fluorescein as indicated in **Figure 5A**. Increased accumulation of sodium fluorescein within the tumor was observed after BIA administration, in comparison with untreated controls (**Figure 5B**). Analysis of the fluorescent signal showed significant accumulation in the tumor, but not in healthy brain, suggesting that BIA administration promoted intra-tumoral uptake of sodium fluorescein.

**Figure 5.**
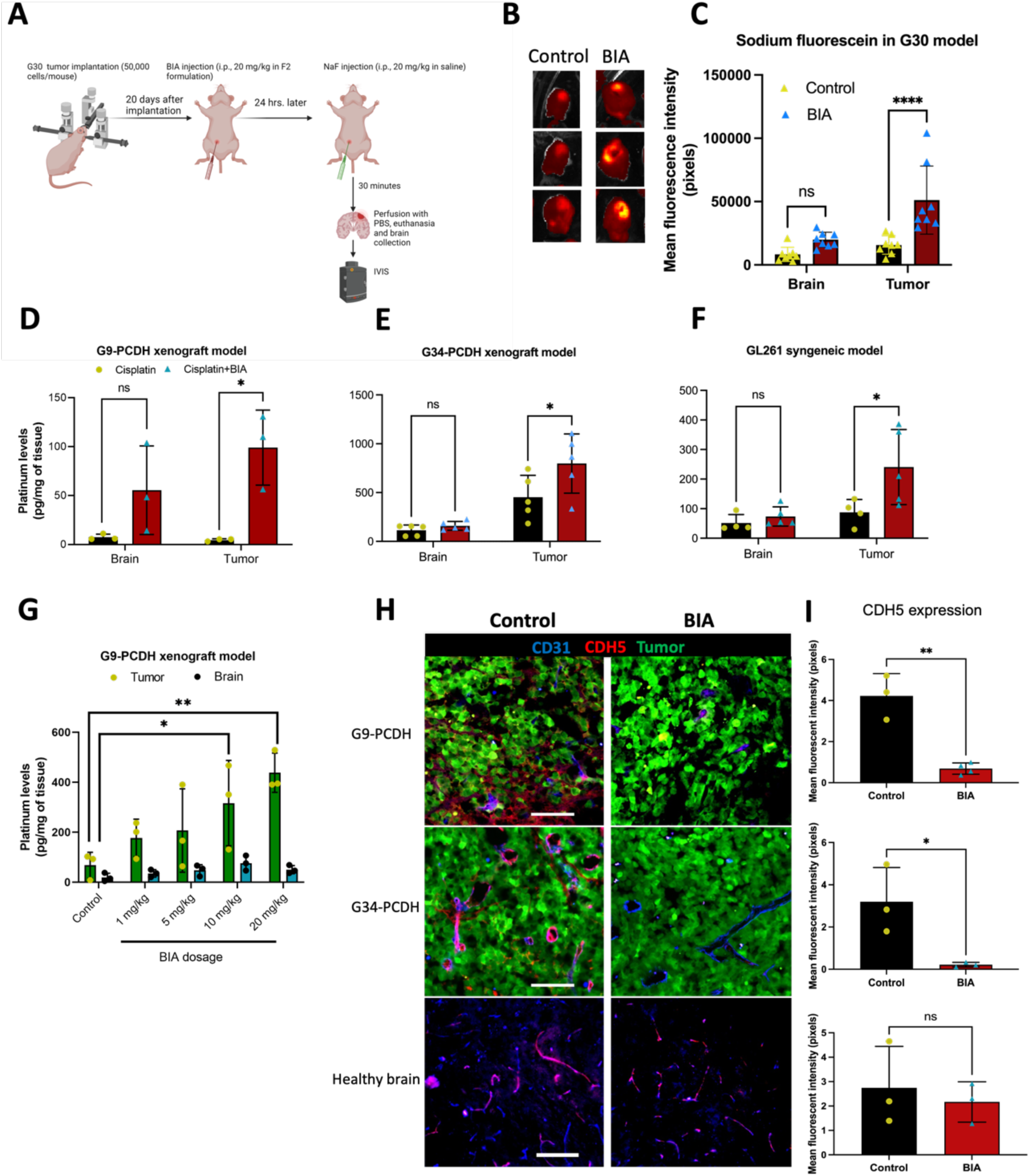
Systemic administration of BIA increases the selective uptake of sodium fluorescein and platinum chemotherapy in murine GBM tumors. **(A)** Workflow schematic of BIA administration and subsequent injection of sodium fluorescein for BTB permeability assessment. **(B)** In vivo imaging system (IVIS) pictures of G30-tumor bearing brains from mice injected with BIA and sodium fluorescein as shown in **(A)**. **(C)** Quantification of image intensity was performed with ImageJ. Mean and standard deviation are shown, n=7-8. Unpaired t-test for statistical significance, ** p=0.0024. (D) Platinum quantification via ICP-MS of brain and tumor tissue from tumor-bearing mice injected with cisplatin in G9-PCDH, **(E)** G34-PCDH and **(F)** GL261 murine models. Cisplatin (5 mg/kg) was administered 24 hours after BIA injection. Mean and standard deviation are shown, n=3-5/group. Two-way ANOVA test was performed, * p<0.05. **(G)** Platinum quantification via ICP-MS of tumor and brain tissue of a G9-PCDH tumor-bearing xenograft model administered with increasing BIA doses. Mean and standard deviation are shown, n=3/group. Two-way ANOVA test, * p=0.0177, ** p=0.0016. **(H)** Immunofluorescence imaging from frozen and sectioned brain tissue from G9-PCDH and G34-PCDH xenograft murine models, 24 hours after injection with 20 mg/kg of BIA. Tumor (green), CDH5 (red) and CD31 (blue) are shown. Scale bars at 100 μm. **(I)** CDH5 fluorescence quantification from experiment in (E) using ImageJ. Mean and standard deviation are shown. Unpaired t-test (n=3/group). ** p=0.0013, * p=0.0334.

We then interrogated whether BIA treatment could increase the intratumoral accumulation of cisplatin, a non-brain penetrant drug. For this, we injected 5 mg/kg of cisplatin and allowed circulation in the system for 5 hours. We collected processed the tissue downstream (see *Materials and methods* section) for Inductively Coupled Mass-Spectrometry (ICP-MS) analysis-based platinum quantification. (**Supplementary Figure 6A**). Pre-treatment with BIA permitted significant cisplatin intra-tumoral accumulation in patient-derived (**Figures 5D and 5E**) and syngeneic murine GBM tumors (**Figure 5F**). Importantly, no significant difference of uptake was seen in contralateral healthy brain regions, indicating that BIA acts selectively in the tumor but not in the brain. Moreover, no difference in platinum accumulation was seen in peripheral tissues such as heart or liver, thereby supporting the notion that BIA selectively increases cisplatin uptake in tumor but not healthy tissue (**Supplementary Figure 6B**).

Further studies showed that the uptake of cisplatin is dependent on the dose of BIA (**Figure 5G**). To test possible mechanisms of how BIA operates in augmenting drug accumulation in tumors, we treated GBM cells (**Supplementary Figure 6C**) and brain endothelial cells (**Supplementary Figure 6D**) with BIA and cisplatin simultaneously. In either case, we did not observe any advantage in drug accumulation due to BIA addition, suggesting that direct cellular internalization is not a mechanism of operation for BIA. In fact, treatment of endothelial cells with BIA did not show any changes of protein levels of CAV1 or MFSD2A (**Supplementary Figure 6E**), important molecular actors in endocytosis and transcytosis in the BBB.

Next, we evaluated CDH5 expression in our patient-derived xenograft GBM models and its potential alterations upon BIA treatment. Administration of BIA showed a striking decrease of CDH5 in CD31+ endothelial cells and in CDH5+ tumor cells 24 hours after treatment (**Figures 5H and 5I**). On the other hand, we did not observe significant changes in expression of CDH5 in contralateral healthy brain regions, which is consistent with the observation that increased drug delivery effects due to BIA are tumor-associated endothelium-specific. Additionally, we assessed the expression of ZO-1 and Claudin-5 in these tissues. We observed mild reductions of ZO-1 expression as well, but no visible differences in Claudin-5 staining (**Supplementary Figure 6F**). Collectively, these data provide evidence that BIA selectively targets the tumoral vasculature at the BTB, which downregulates CDH5 expression, disrupting tight junction formation and increasing accumulation of chemotherapy in murine GBM tumors.

### BIA potentiates cisplatin cytotoxicity by fostering its DNA-damage capacity in GBM cells

Prior to animal efficacy studies, we also asked whether BIA and cisplatin in combination could also show a therapeutic advantage than administration of either agent alone. Several studies have shown cytotoxic synergy of small-molecule kinase inhibitors in combination with cisplatin in cancer ^26–28^. Accordingly, we cultured a panel of patient-derived GBM neurospheres and treated with BIA and cisplatin combination, with single-treatment groups as controls (**Figure 6A**). Using a cell viability assay, we observed that combination of BIA dramatically increased the cytotoxic effects of cisplatin alone. The most significant combinatorial effects were observed at cisplatin concentrations of 1 μM and below. BIA single-treatment controls only mildly reduced cellular ATP production. In accordance, the BIA/cisplatin combination decreased the neurosphere formation capacity and growth of G9 and G34 cells (**Supplementary Figures 7A-7D**). To identify potential synergistic interactions between BIA and cisplatin, we utilized SynergyFinder 3.0 software. BIA potentiated cisplatin toxicity (overall δ-score=8.24), at a concentration of 2.5 μM and below (**Figure 6B**). We also identified a high likelihood of synergy (highlighted area, δ-scores >10) at the lower doses for cisplatin (∼0.6 μM −2.5 μM) in combination with all tested BIA doses (**Figure 6C**). Interestingly, at the upper cisplatin dose-ranges, its interaction with BIA remained non-synergistic. Thus, cisplatin and BIA in combination show synergistic anti-glioma cytotoxic effects.

**Figure 6.**
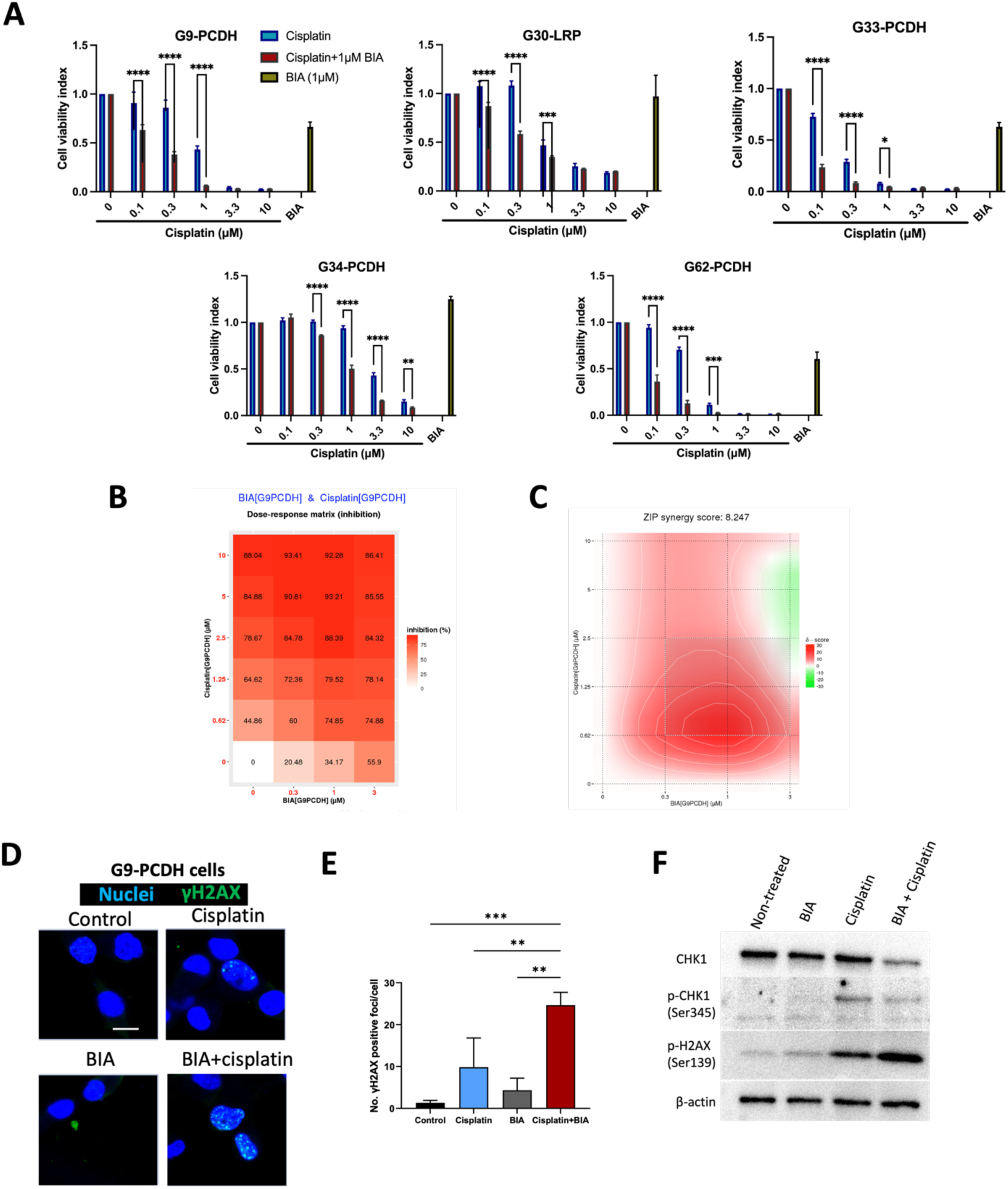
BIA potentiates platinum-based cytotoxicity by targeting DNA-repair pathways in patient-derived GBM cells. (**A**) GBM cell viability of BIA and cisplatin combination treatment. Cisplatin doses are indicated in x-axis, BIA remained at a constant concentration of 1 μM (5 days exposure). Mean and standard deviation are shown, n=3/group. (**B**) Dose-response matrix showing inhibition percentage of BIA and cisplatin combinations at various concentrations for 5 days using SynergyFinder 3.0. G9-PCDH cells were treated, and viability analyzed as indicated in the cell viability assay section (see Materials and Methods). (**C**) ZIP method synergy score of BIA and cisplatin combinations. The overall average ***δ***-score is indicated on top of the chart. The dose combinations showing an increased likelihood of synergy are highlighted. (**D**) Immunofluorescence staining of γH2AX (green) and nuclei (blue) in G9-PCDH cells treated with 1 μM of cisplatin and/or BIA, for 72 hours. Representative image of 3 pictures per condition. Pictures taken at 40x, scale bar=20 μm. (**E**) Quantification of γH2AX foci from (**D**) using ImageJ. (**F**) Western blot of G9-PCDH cells treated with 1 μM of cisplatin and/or BIA, for 72 hours, probing for the p-CHK1 (Ser345), total CHK1 and H2AX (Ser139) proteins. GAPDH was used as loading control. Representative image from triplicate experiments.

Next, we assessed the DNA damage levels of the BIA/cisplatin combination by IF imaging of γH2AX nuclear foci. This showed that the BIA/cisplatin combination significantly augmented the frequency of γH2AX foci in the nucleus of GBM cells above single-treatments and non-treated controls (**Figures 6D** and **6E**). This increase in γH2AX events in the BIA/cisplatin combination was also observed by flow cytometry, which correlated with loss of cell cycle progression (**Supplementary Figures 7E and 7F**). We evaluated the phosphorylation and protein levels of CHK1, an important regulator of the DNA damage response during cisplatin exposure ^29^. Simultaneous exposure of BIA and cisplatin reduced the expression of CHK1 and its activation (Ser345) greater than single-treatment controls. In turn, γH2AX levels were induced upon this combination (**Figure 6F**). Given the strong depletion of CHK1 activity, we performed siRNA-dependent knock-down in our GBM cell lines. Use of siCHK1 increased the susceptibility of these cells to cisplatin titrations, mainly at concentrations below 1 μM (**Supplementary Figure 7G**), supporting the notion that targeting of CHK1 is an important factor in the BIA-induced potentiation of cisplatin cytotoxicity.

### Administration of BIA prior to cisplatin dosing shows pre-clinical efficacy in patient-derived xenografts of GBM

Finally, we investigated whether BIA and cisplatin combination regimens could provide a therapeutic effect in our intra-cranial GBM murine models. We proceeded with a dose-regime of BIA pre-administration 24 hours before cisplatin injection at 5 mg/kg to promote and maintain increased platinum delivery (**Figure 7A**). The BIA and cisplatin combination regimens prolonged the survival of tumor-bearing mice significantly (p = 0.0052) over the single-treatment and control arms, indicating efficacious results by this approach (**Figure 7B**).

**Figure 7.**
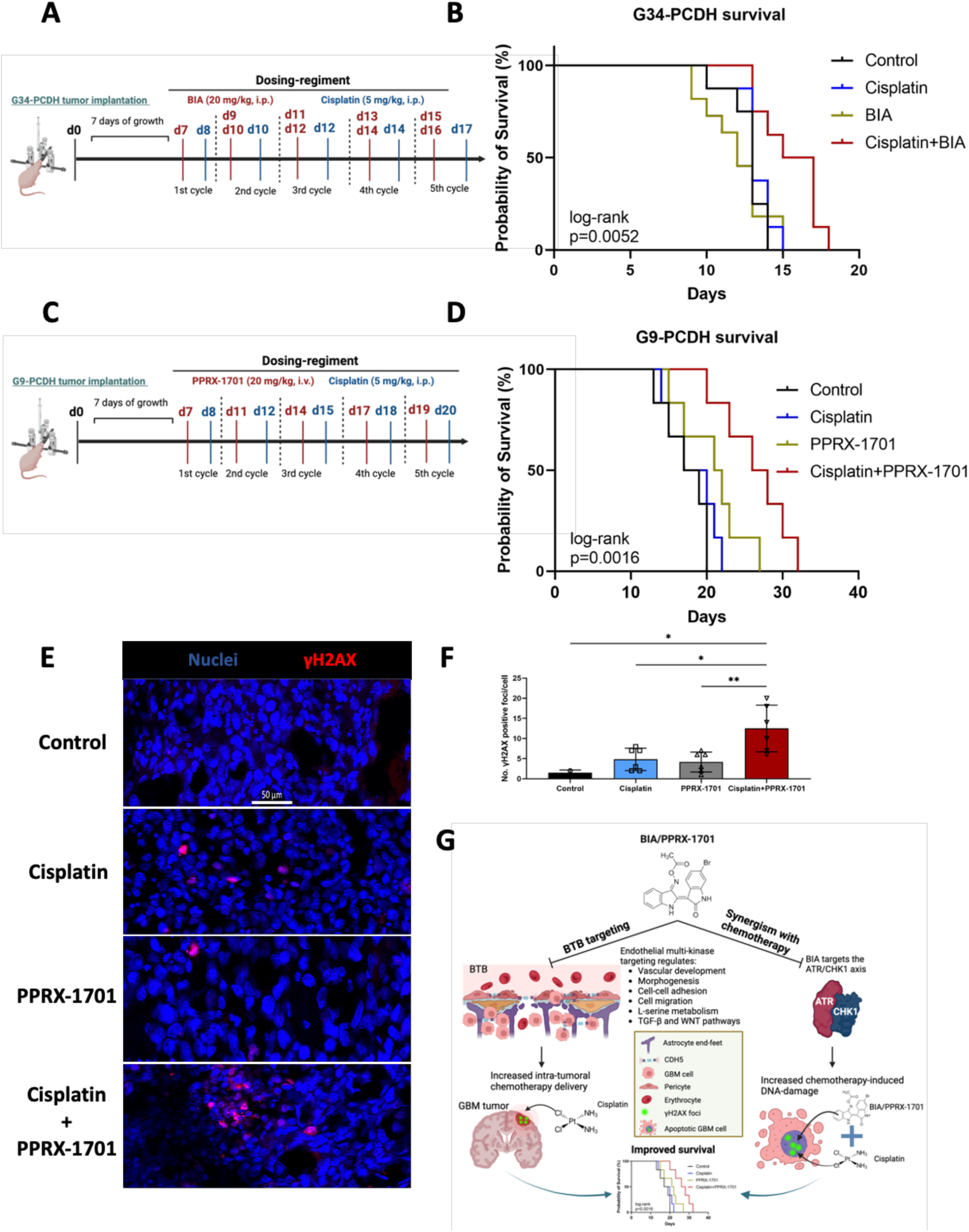
Systemic administration of BIA in combination with cisplatin treatment shows enhanced pre-clinical efficacy in murine GBM models. (**A**) and (**C**) Diagrams of the experimental design for G34-PCDH and G9-PCDH xenograft efficacy studies using BIA/PPRX-1701 and cisplatin combinations. (**B**) Efficacy studies of G34-PCDH xenograft using BIA and (**D**) PPRX-1701 in combination with cisplatin. For PPRX-1701 studies, vehicle formulation was used as control and in combination with cisplatin. n=8/group. Log-rank test analysis for statistical significance. (**E**) Confocal immunofluorescence imaging of γH2AX (Alexa Fluor 647, red) nuclear foci from tumor tissue collected from study (D). Nuclei were stained with Hoechst 33342 (blue). Representative pictures taken at 20x. Scale bar=50 μm. (**F**) Quantification of γH2AX foci from (**E**) using Image J, n=6/group. Ordinary One-way ANOVA was performed for statistical evaluation. * p=0.01, ** p=0,0086. (**G**) Schematic of proposed model of BIA/PPRX-1701 mechanism of action and its effects in GBM tumor drug delivery and efficacy.

BIA is highly hydrophobic, making it difficult to dissolve in physiological solutions, which limits its clinical translation. To address this, we used PPRX-1701, a formulation of BIA, designed for improved *in vivo* delivery ^30^, which inhibits GSK3 as indicated by a G9-TCF cell line reporter (**Supplementary Figure 8**). Previously, we have shown that PPRX-1701 is not toxic when administered systemically in C57/BL6 mice, as shown by liver and spleen histology ^30^. We implanted a second patient-derived GBM xenograft model (**Figure 7C**) and performed systemic pre-administrations of PPRX-1701 before cisplatin injections. Combination of PPRX-1701 with cisplatin was also more efficacious in comparison with vehicle control plus cisplatin, PPRX-1701 alone and non-treated controls (p = 0.0016) (**Figure 7D**). Assessment of DNA damage by γH2AX staining indicated that PPRX-1701 enhanced the genotoxicity of cisplatin, correlating with the extended survival observed (**Figures 7E and 7F**).

Altogether, our data highlights potential molecular targets associated to the BTB in GBM. In addition, we demonstrated that BIA exerts pre-clinical efficacy in GBM murine models through its dual capacity to selectively target transcriptional programs of the BTB, promoting intra-tumoral drug delivery and by showing cytotoxic synergistic effects with DNA-damaging chemotherapy (**Figure 7G**).

## Discussion

Effective drug delivery remains a major barrier for the treatment of brain tumors. Here, we have identified a network of genes associated with the BTB in GBM and have demonstrated the dual functionality of the indirubin-derivative, BIA, to increase intra-tumoral drug delivery by targeting the BTB-associated gene network and enhance chemotherapy cytotoxicity via DNA-repair machinery modulation. Our work should provide grounds to establish further studies for progressing BTB-targeting approaches towards clinical application.

We used an *in silico* strategy to identify a set of 12 genes (BTB-genes) with elevated regional expression within the tumoral endothelium in GBM. Moreover, our spatial transcriptomic data analysis showed high spatially-clustered expression of these genes in GBM clinical samples, suggesting a functional relevance for this disease. Most of these genes have been associated with angiogenesis and blood-vessel recruitment, especially *ACVRL1* (also called *ALK1*), *CD93*, *ENG*, *FLT4* (VEGFR3) and *PDGFRB*. Previous studies ^31^ have indicated the co-expression of *ACVRL1*, *CDH5*, *CLEC14A*, *PECAM1*, *ENG*, *GRP4*, *ROBO4* and *PCDH12* in the tumor-associated endothelium in several solid tumor types, including GBM. In fact, this gene set has been related to vascular development, blood vessel morphogenesis and tumor angiogenesis processes. These pathways have also been identified in endothelium of primary GBM specimens ^32,33^. The mentioned reports support our findings and indicate the functional relevance of these genes and vascular developmental processes to the BTB in GBM. Our work links the modulation of these BTB-genes and vascular pathways to alter BTB permeability for improved drug delivery in tumors. Future work by us would involve functional studies on these molecules for deeper understanding of their involvement in BTB biology.

Our screening led us to identify CDH5 (VE-cadherin) as a central element in the tumoral vasculature transcriptome. CDH5 is fundamental for endothelial barrier integrity, but its role in BTB permeability is not fully understood. CDH5 showed prominent expression in vascular hyperproliferative regions of GBM clinical samples, and its expression was correlated with endothelial markers above non-tumor cortex when analyzed by spatial transcriptomics. CDH5 downregulation strongly correlated with increased drug accumulation after BIA injection in our GBM murine models. Functionally, CDH5 has been associated with enabling vascular mimicry capacity in GSCs ^34^, permitting formation of vascular-like structures that supply with nutrients and facilitate anti-angiogenic therapy resistance. Our findings show that BIA targets CDH5 gene expression in endothelial and tumoral compartments. It is possible that this simultaneous cellular targeting is at least partially responsible for the increased sodium fluorescein and cisplatin accumulation we observed. On the other hand, we encountered altered expression levels of genes involved in TGF-β and WNT signaling pathways. The TGF-β pathway maintains BBB integrity through crosstalk with oligodendrocytes, pericytes and endothelial cells ^35,36^. We observed downregulation of several members of this pathway. The WNT/β-catenin pathway is fundamental for brain and retinal barriergenesis and maintenance, especially the Norrin/WNT7A/B axis ^37,38^. We observed a decrease of *WNT7B*, and WNT ligand receptors *FZD4* and *FZD7*, with simultaneous increase of expression of *WNT4*, *WNT10A* and *WNT11* ligands. Additionally, transcriptional alteration of genes involved in angiogenesis (i.e. *ANGPT2*, *ENG*, *ANG*) and BTB permeability (*S1PR1* and *S1PR3*) was also seen. As such, CDH5 downregulation and alteration of BBB-integrity components might work together to contribute to the BTB permeability modulation exerted by BIA. Future work by our team will focus on functional interrogation of the potential roles of CDH5 in the tumor-associated vasculature and relevance in the permeability of the BTB for drug delivery purposes.

The administration of BIA to tumor-bearing xenograft and syngeneic mice enhanced the accumulation of cisplatin and sodium fluorescein in brain tumor tissue but not healthy brain. The specificity of this effect towards tumorigenic regions remains under study by us. It is likely that BIA, being a small-molecule kinase inhibitor, targets cells with elevated kinase signaling activity, such as the case of angiogenic/proliferative endothelium, but spares slow cycling/quiescent cells that constitute the non-tumorigenic brain vascular networks. Vascular development and motility programs are active in angiogenic endothelial cells, and the multi-targeting quality of BIA can dysregulate multiple elements involved in these pathways. We also observed inactivation by dephosphorylation of the endothelial nitric oxide synthase (eNOS), important blood-pressure regulator, and the p38α/CREB axis, which can control gene expression of CDH5 and other genes important to endothelial biology. The p53 tumor suppressor protein also showed an increase of activating phosphorylation sites (S46) and loss of phosphorylation modulating proapoptotic (S392) and gene regulation (S15) activities. The p53 factor has been involved in modulating endothelial vasodilation and functions in vascular remodeling ^39–41^. On the other hand, BIA promoted the expression of genes relevant to L-serine metabolism and amino-acid transport processes. L-serine has been reported to improve cerebral blood-flow, which provides neuroprotection during CNS disease ^42^. In this regard, normalized blood-flow can also promote drug accumulation in solid tumors ^43,44^. The mTORC1 complex is an important amino-acid sensor, which regulates protein synthesis and energy modulation. We observed an increased phosphorylation of p70 S6 kinase, a downstream target of the mTORC1 pathway. This is consistent with the observation that BIA promotes expression of genes related to amino acid transport and synthesis. This could potentially be result of indirubin-derived metabolic secondary effects, future work should be performed to confirm and address the therapeutic relevance of such observations.

Simultaneous exposure to BIA and cisplatin had a synergistic killing effect in GSC-like cells. This correlated with increased DNA damage and CHK1 inhibition. Other studies have shown indirubin derivatives induce DNA-damage in HCT-116 cancer cells ^45^. However, the present work reveals a novel applicability of BIA, and potentially other indirubins, in combinatorial regimens to synergize with DNA-damaging chemotherapy. Administration of BIA or PPRX-1701 nanoparticles, followed by cisplatin after 24 hours, permitted an extension of survival of two different GBM xenografts. Most likely this improved pre-clinical efficacy stems from the increased platinum delivery intratumorally and the additive cytotoxicity exerted by both agents. Given this finding, other DNA-damaging chemotherapeutics should be screened in combination with BIA to identify alternative drug candidates that would benefit from the increased accumulation and BIA anti-neoplastic synergism in GBM treatment. The mechanism of how BIA downregulates CHK1 expression at the protein level, and what alternative chemotherapy modalities will benefit with BIA remains in ongoing study by us.

Altogether, our work reveals novel molecular markers of the BTB, which in future studies should be functionally characterized to understand their role in the biology of the BTB-GBM interaction. The identification of BIA as a selective regulator of BTB permeability for improved drug delivery and potentiating agent of DNA-damaging chemotherapy supports the use of BIA in further pre-clinical and clinical studies of GBM. Primarily, further research should be pursued on screening for non-BBB penetrant chemotherapies and biologicals that would benefit from higher intra-tumoral internalization in combination with BIA, such as small molecule inhibitors, chemotherapies, and therapeutic antibodies.

## Materials and methods

### BTB-gene in silico screening

To identify genes related to BTB function, we initiated an *in silico*-based approach by accessing bulk RNA-sequencing data from The Cancer Genome Atlas via the cBIO portal for Cancer Genomics (https://www.cbioportal.org/). We initiated a correlation analysis of genes co-expressed with endothelial markers PECAM-1 (CD31), Von-Willebrand Factor (VWF), C-lectin 14 type A (CLEC14A) and CD34, previously identified as useful markers of GBM vasculature ^17^. A selection of top-50 genes (Spearman’s rank correlation) commonly observed in 3 out of the 4 markers was done and interrogated their expression levels in GBM tumors in comparison to healthy brain by using the GlioVIS portal (http://gliovis.bioinfo.cnio.es/) to visualize the Rembrandt study ^46^. Those genes significantly elevated in tumor over healthy brain were selected as candidate BTB targets due to their possible relevance in GBM. Then, their regional expression in GBM was assessed by using the IVY GAP (https://glioblastoma.alleninstitute.org/) data visualized in the GlioVIS portal. Using this tool we confirmed their expression in microvascular proliferative regions, which are associated with the vasculature in tumors. All graphs of GlioVIS and cBIO portal datasets were generated in the corresponding websites and pairwise t-tests performed for statistical significance test.

### Gene Ontology analysis

For Gene Ontology (GO) analyses, we used the EnrichR (https://maayanlab.cloud/Enrichr/) website, generated by the Ma’ayan’s lab ^47–49^. We used the GO Biological Process 2023 visualization tool to identify biological processes of the identified gene-sets. The Appyters notebook ^50^ linked to EnrichR was used for graphics visualization.

### Gene interaction network analysis and gene-set clustering

Gene-sets were submitted to the Search Tool for the Retrieval of Interacting Genes (STRING, https://string-db.org/ ^51^). Scores were set to medium interaction (0.4). For interaction analysis of the genes targeted by BIA, we selected genes upregulated and downregulated by BIA equals or above 2-fold change (Log2). Only genes that presented interaction were associated by a 4-kmeans clustering. Gene-sets comprising each cluster were submitted to GO analysis (using EnrichR as mentioned above) and ranked by p-value significance. The most significant pathway by this method is indicated by color-code in each cluster.

### Spatial transcriptomics data set analysis and clustering methods

In this paper, four specific datasets out of a larger set of 28 were focused on data available from Ravi *et al*., 2023 ^18^, dataset that was deposited in Datadryad (https://doi.org/10.5061/dryad.h70rxwdmj) by the authors. These datasets were collected from patients with tumor and cortex control samples. To analyze the effect of CDH5, we started clustering our spatial dataset and visualizing the gene expression spot information with spatial dimensions using the SPATA2 package in R-studio ^52^ (https://github.com/theMILOlab/SPATA2). Additionally, utilizing the Seurat package (v5.0.0) in R (v4.2.2), the spatial transcriptomics data were processed in several steps. Initially, the data were loaded and preprocessed, followed by normalization using the Log Normalization method. Variable features were identified, and the data were scaled accordingly. Principal Component Analysis (PCA) was then applied to reduce the dimensionality of the dataset, with emphasis placed on the top 20 principal components for subsequent cluster and neighbor analysis based on PCA dimensions. The data were visualized in two dimensions using Uniform Manifold Approximation and Projection (UMAP). The tumor and cortex control datasets were merged into a single Seurat object using the merge function (Seurat::merge()). Subsequently, the spatial layers were processed to facilitate visualization of the data in two dimensions. Gene expression patterns were analyzed using the same dimension reduction plot, and expression levels were assessed with violin plots within each cluster identified by the Seurat algorithm.

### Spatial Gene Ontology Analysis

The Gene Ontology (GO) analysis employed a cluster-based methodology conducted in R, with clusters determined by the Seurat algorithm. Initially, differential expression analysis was conducted in Seurat to identify genes and their associated cluster information within the samples (Seurat::FindAllMarkers()). Subsequently, genes were individually grouped based on their clusters, and GO analysis was performed using the enrichGO function in the clusterProfiler package (https://bioconductor.org/packages/release/bioc/html/clusterProfiler.html). A reference genome-wide annotation for human, primarily utilizing mapping via Entrez Gene identifiers, was obtained from the org.Hs.eg.db package within the Bioconductor library and converted into a data frame. Visualization of the GO data was accomplished using the GOplot package (GOplot::GoBubble()). For optimal visualization, only the cluster containing CDH5 was selected and depicted in the bubble plot.

### Spatial Pathway Analysis

Pathway analysis was conducted using a gene-based approach, where signature genes corresponding to each pathway were sourced from the MsigDB (https://www.gsea-msigdb.org/gsea/msigdb/human/genesets.jsp). Specifically, our focus was on the WNT pathway, Vegf angiogenesis pathway, and TGFB pathway, with gene extraction performed using the msigdb package (https://www.bioconductor.org/packages/release/data/experiment/html/msigdb.html). These pathways are categorized within the Curated Gene Sets collection (C2 gene sets) under the Biocarta sub-collection. The expression patterns of individual genes derived from this methodology were visualized using the FeaturePlot function in R, enabling two-dimensional visualization.

### Mice

Female Nu/Nu mice (Envigo) and C57/BL6 (Charles River Laboratories) aged 8 weeks were used for *in vivo* experiments. All our procedures followed the guidelines by the Institutional Animal Care and Use Committee (IACUC) with support of the Center for Animal Resources and Education (CARE) at Brown University.

### Cell lines

Glioma-stem cell-like cell lines G9-PCDH, G34-PCDH, G33-PCDH, G62-PCDH and G30-LRP were obtained and cultured as previously described ^12,30,53^. Briefly, cells were grown as neuro-spheres using Neurobasal medium (Gibco) supplemented with 20 ng/ml of human recombinant EGF (Peprotech), 20 ng/ml of human recombinant FGF (Peprotech), 2% B-27 supplement (Thermo Fisher Scientific), 0.1% GlutaMax (Thermo Fisher Scientific), and 0.1% penicillin/streptomycin (Thermo Fisher Scientific). Cells were left to grown at least overnight for sphere formation. For single-cell dissociation, Accutase (Gibco) was used for 5 minutes at 37 degrees. For culturing GL261-Luc2 cells, we used 10% fetal bovine serum (Gibco), with 0.1% GlutaMax and 0.1% penicillin/streptomycin in DMEM/F12 media (Gibco).

Growth and culturing of immortalized human cerebro-microvascular endothelial cells (HCMEC/D3) (Sigma), primary human brain microvascular endothelial cells (HBMEC) (ScienCell), human primary astrocytes (Lonza Biosciences) and human primary pericytes (ScienCell) was performed as previously reported ^54,55^. Briefly, HCMEC/D3 and HBMEC cells were cultured in Endothelial Cell Media (ScienCell) supplemented with fetal bovine serum, endothelial cell growth supplement and penicillin/streptomycin as provided by the company. Astrocyte and Pericyte cells were grown in complete formulations of astrocyte cell media (ScienCell) and pericyte cell media (ScienCell), respectively. For immunostaining experiments HCMEC/D3 and HBMEC cells were grown in type 1 rat collagen-coated plates. These endothelial cells were used below passage 20 for maintenance of their BBB properties.

### Cell viability assay

Cells were plated at a density of 1500 cells/well in black-well clear bottom 96-well plates and left growing in culture conditions overnight. Next day, cells were treated with titrating doses of the indicated compounds. For BIA only cytotoxicity studies, cells were incubated with BIA for 96 hours. For BIA and cisplatin combinatorial studies, cells were incubated with BIA and cisplatin, and corresponding controls, for 5 days. Next, we used the Cell-Titer Glo 3D (Promega) following provider’s guidelines and quantified for luminescence signal using a Molecular Devices SpectraMax M2 plate reader. Conditions were repeated in triplicates.

### Growth in Low Attachment (GILA) assay

Fluorescently labelled GBM cells (G9-PCDH and G34-PCDH, GFP-labelled) were plated in clear ultra-low attachment 96-well plates (Costar) with a density of 2000 cells/well using 100 μl of complete Neurobasal medium. Then, cells were centrifuged at 1200 rpm for 3 minutes. Cells were treated as indicated above and fluorescence visualized using a Nikon Eclipse Ti2 microscope. Sphere diameter was measured using Image J software. Conditions were repeated in triplicates.

### Synergy analysis of BIA and cisplatin combinations in GBM neuro-spheres in vitro

To identify if the BIA and cisplatin combinations present synergistic anti-neoplastic effects in GBM cell line neuro-spheres, we used the SynergyFinder 3.0 software ^56^. For this, cell viability assays (see above) were performed. Concentrations of 0 μM, 0.3 μM, 1 μM and 3 μM of BIA were added in combination with 0 μM, 0.62 μM, 1.25 μM, 2.5 μM, 5 μM and 10 μM of cisplatin, accordingly, for an exposure duration of 5 days. Cell Titer Glo 3D assays were performed for cell viability assessment. SynergyFinder 3.0 analysis was done with LL4 curve fitting, with outlier correction, following a ZIP synergy score. We performed a ZIP-based analysis since this model low false-positive rates while calculating synergy of anti-oncogenic drugs ^57^. For reference, a δ-score of less than −10, could signify antagonism, −10 to 10 could signify additivity, and above 10 could signify synergism.

### RNA-sequencing of HCMEC/D3 cells treated with BIA

For RNA-sequencing, HCMEC/D3 cells were plated at a density of 500,000 cells/well in a 6-well plate. Left to grow for 24 hours, and then treated with 1 μM of BIA or DMSO (control). After 24 hours, cells were collected and processed for RNA extraction using the column-based RNeasy kit (QIAGEN), following provider’s instructions. RNA quality and quantity were quantified using a Nanodrop™ One (Invitrogen). At least 500ng of RNA was submitted for bulk RNA-sequencing at GeneWiz (Azenta Life Sciences). QC was accessed, and library was prepared with Poly(A) selection. Sequencing was performed using Illumina HiSeq.

Differential gene expression on the RNA-seq raw data (FASTQ files) was analyzed by Azenta Life Sciences using DESeq2 aligning to human transcriptome. Data QC was verified. Log_2_ fold change (Log2FC) was calculated by Log_2_ (BIA group mean normalized counts/Control group mean normalized counts). The Wald test p-value and Benjamini-Hochberg adjusted p-value were calculated. A heatmap and volcano plot of top adjusted p-value differentially expressed genes (DEGs) in ensemble ID annotation bi-clustering to treatment conditions were generated. Control and BIA groups consisted of three-independent samples.

### Immunofluorescent (IF) staining

For IF staining of HCMEC/D3 endothelial cells, we coated 8-well Nunc™ Lab-Tek™ chamber slides (Thermo Fisher Scientific) with 1X Type 1 rat-tail collagen (Corning) following provider’s instructions. Then, we plated at a density of 50,000 cells/well and left in culture for 72 hours to allow for barrier formation. Next, we treated with BIA or control for 24-48 hours. Cells were then fixed with 10% formalin (Thermo Fisher Scientific) for 10 minutes, permeabilized for 30 minutes using 0.01% Triton X-100 and blocked with 0.1% normal donkey serum (Calbiochem) for 1 hour in 0.025% Tween-20 (Thermo Fisher Scientific) in Phosphate-Buffered Saline (PBS) (Gibco). Then, primary antibodies were added: mouse anti-CDH5 (VE-cadherin, BioLegend) 1:100, rabbit anti-Claudin-5 (Thermo Fisher Scientific) 1:100, and mouse anti-ZO-1 (Invitrogen) 1:100, and incubated overnight in the cold. Next day, secondary antibodies were used for 2 hours at room termperature: Alexa Fluor 594 anti-mouse (1:500), Alexa Fluor 594 anti-rabbit (1:500), and Alexa Fluor 647 anti-mouse (1:500), all of these Thermo Fisher Scientific. For cytoskeleton staining Phalloidin-iFluor 488 (Abcam) 1:1000 for 30 minutes and nuclei staining using Hoechst 33342 (Thermo Fisher Scientific) 1:1000 for 5 minutes, at room temperature.

For GBM cell staining, cells were cultured in 10% DMSO in complete Neurobasal media for 2 days, and then plated at a density of 50,000 cells/well in 8-well Nunc™ Lab-Tek™ chamber slides. IF staining was performed as indicated above for endothelial cells. Primary antibodies used: Rabbit anti-γH2AX (Ser139) (Cell Signaling) at 1:100 dilution. A goat anti-rabbit Alexa Fluor 647 was used at 1:500.

For mouse brain tissue staining, brains were collected from CO_2_ euthanized and PBS perfused tumor-bearing mice, and fixed in 10% formalin for 72 hours on rotation in the cold. Then, brains were transferred to 30% sucrose for 3 days at 4 degrees Celsius under rotation. Before cryosectioning, brains were frozen at −80 degree Celsius for more than 30 minutes, embedded in Optimal Cutting Temperature (OCT) compound (Fisher) and transferred to −27 degrees Celsius to a cryostat (Leica CM1950) for sectioning (20 μm thickness). Sections were placed on slides and staining followed as indicated above. All pictures were taken using a LSM 880 Zeiss confocal microscope.

### BBB spheroids and dextran permeability assay

BBB spheroids were grown and cultured with a FITC-conjugated (70kDa) fluorescent dextran (Millipore Sigma) as previously reported ^21,58,59^. BBB spheroids were grown for 48 hours and then treated with BIA at increasing doses for 72 hours. Then, spheroids were collected and stained as indicated above for CDH5 and F-actin (Phalloidin). In the case of fluorescent dextran incubation, BBB spheroids were collected in an 1.5 ml microtubes (Eppendorf) and incubated for 3 hours at 37 degrees Celsius. Pictures were taken by confocal microscopy. For dextran permeability measurement, 21 images using Z-stack layers of 5 μm intervals for achieving a total depth of 100 μm within the sphere. Fluorescent-dextran intensity from maximal intensity projection was quantified using Image J (National Institute of Health).

### Trans-endothelial electrical resistance

HCMEC/D3 or HBMEC cells were plated in 8W10E+ PET 8-well arrays (Applied Biophysics) at a density of 100,000 cells/well in 500 μl. These arrays were placed in a pre-stabilized ECIS Z-Theta instrument (Appled Biophysics). Using the ECIS Z-Theta software (Applied Biophysics), measurements were set to 4000 and 64000 Hz every 30 minutes. Cells were left to grow and form a barrier for 48-72 hours (normally, a resistance plateau would be reached, and capacitance showed at ∼10nF for 64000 Hz). Cells would then be treated with BIA and left to grow up to 5 days, with frequent drug-containing media re-addition for maintenance of the culture. Resistance (Ω) and capacitance (nF) were recorded and plotted.

### Real-time PCR

Total RNA from GBM and HCMEC/D3 cells was obtained and processed as indicated above. For cDNA generation, we used 1 μg of RNA and processed with with the iScript™ cDNA synthesis kit (BioRad), following the protocol indicated by the provider. All primers were designed using NCBI Primer-Blast tool. Detailed information on primer sequence can be found in Supplementary **Table 3**. Gene expression levels were quantified using PowerUp SYBR Green Master Mix (Applied Biosciences) on QuantStudio 6 Pro System (Applied Biosciences), normalized by housekeeping gene GAPDH expression and represented as relative expression using comparative ΔΔC_T_ method.

### Western blot

HCMEC/D3 and GBM cell lysates were collected in RIPA Buffer (Thermo Fisher Scientific) supplemented with 1x protease/phosphatase inhibitor cocktail (Cell Signaling). Lysate collection from murine tumor tissue samples (∼ 30mg) was performed under homogenization using 23G and 26G needles. Total protein concentration was measured using Pierce 660nm Protein Assay Reagent (Thermo Fisher Scientific) at 660nm absorbance in Molecular Devices SpectraMax M2 plate reader. Samples were incubated in 1x Laemmli sample buffer (BioRad) at 95 degrees Celsius for 5 minutes before loading onto 10% Mini-PROTEAN TGX precast protein gel (BioRad). PageRuler Plus Pre-stained Protein Ladder (Thermo Fisher Scientific) was used as ladder. Blocking was performed in 5% milk with 0.1% Triton X-100 in 1x PBS (TBST) (Gibco) for 1 hour at room temperature under shaking. Primary antibodies used were incubated in the cold under shaking overnight: anti-pCHK1 (Ser345) (Cell Signaling Technologies) 1:100, anti-CHK1 (Cell Signaling Technologies) 1:1000, anti-pH2AX (Ser139) (Cell Signaling Technologies) 1:1000, anti-MFSD2A (Proteintech) 1:500, anti-CAV1 (Proteintech) 1:1000, anti-CD144 (VE-cadherin) (Thermo Fisher Scientific) 1:1000, anti-β-actin (Cell Signaling Technologies) 1:2000. Appropriate secondary antibodies, Goat anti-mouse-HRP (Sigma) or Goat anti-rabbit-HRP (Sigma) in 5% milk in 1x TBST with 1:5000 dilution for 1 hour at room temperature.

### Phospho-kinase array

HCMEC/D3 cells were plated at a density of 1 million cells and treated with either 1µM BIA or vehicle DMSO for 24hr. Cell lysates were collected with manufacturer provided Lysis Buffer 6 supplemented with 10 µg/mL Aprotinin (Tocris), 10µg/mL Leupeptin hemisulfate (Tocris) and 10µg/mL Pepstatin A (Tocris) for protein preservation. 50µg of lysate from each sample were loaded into each membrane. All experiment procedures were performed using the Proteome Profiler Human Phospho-Kinase Array Kit (R&D Systems) following manufacturer’s protocol.

### siRNA transfections

G9-PCDH and G30 cells were cultured to approximately 60% confluency and transfected using Lipofectamine RNAiMax Transfection Reagent (Invitrogen) for 1 day, and then replated for western blot or cell viability assays. All experimental steps followed manufacturer’s protocol. siCHK1 (Ambion) was used for CHK1 depletion. MISSION siRNA universal negative control (Sigma-Aldrich) was used as control siRNA.

### Flow cytometry for cell cycle, DNA damage and apoptosis assays

For cell cycle analysis, 100,000 HCMEC/D3 cells were plated in 6-well plates and treated with indicated concentrations of BIA or control for 48 hours. Then, cells were washed twice with PBS and fixed/permeabilized with 5 ml of cold 70% Ethanol added dropped-wise while vortexing at low speed. Cells were stored for 1 day at −20 degrees Celsius, washed three times with PBS and treated with 20 µg/ml RNAse I (Thermo Fisher Scientific) and stained with anti-Ki-67 FITC-conjugated (1:1000) (BD Biosystems) and 1.5 µM propidium iodide (Thermo Fisher Scientific). After 30 minutes of incubation in the dark, cells were analyzed using a CytoFLEX system (Beckman Coulter). 50,000 events were counted, and data was analyzed using the CytoFLEX system software (Beckman Coulter).

For apoptosis assessment, HCMEC/D3 cells were treated as indicated above for 72 hours with BIA at indicated doses. Cells were collected from the 6-well plates and washed three times with PBS. Then, incubated with SYTOX™ Blue nucleic acid stain (5 mM) with a dilution of 1:1000 for 15 minutes. Cells were submitted and analyzed in the CytoFLEX system and its software as indicated above.

For DNA damage and cell cycle assessment of BIA and cisplatin, G62 cells were plated at a density of 100,000 cells/well in a 6-well plate and grown in complete Neurobasal media. Cells then were treated with BIA and/or cisplatin and control for 72 hours. Next, cells were collected and washed three times with PBS and stained with 1:500 of FITC anti-γH2AX Phospho (Ser139) (BioLegend) antibody and Propidium iodide (1 mg/ml) at 1:1000 dilution for 30 minutes in the dark. Cells were taken for analysis in a BD Fortessa cytometer and data analyzed using a FloJo software (BD Biosciences).

### Cell counts

HCMEC/D3 cells were counted and plated at a density of 300,000 cells/well in a 6-well plate. After overnight growth in culture conditions, DMSO or BIA were added at 1 μM or 5 μM. Cells were counted every 2 days and media with fresh BIA or DMSO replaced for continuous growth.

### Secretome quantification

For cytokine analysis of brain endothelial cells after BIA exposure, HCMEC/D3 cells were plated at a density of 500, 000 cells/well in a 6-well plate. Cells were treated with indicated doses of BIA or DMSO for 48 hours. Then, 1 ml of media was collected and processed for cytokine quantification in a Luminex platform (Thermo Fischer Scientific) following the provider’s instructions.

### BIA and PPRX-1701 preparation

BIA powder stocks (Millipore Sigma) were resuspended in DMSO at a concentration of 10 mM (*in vitro* usage) or 100 mM (*in vivo* usage). For animal experiments, 100 mM BIA was dissolved in 2% Tween-20 (Thermo Fisher Scientific), 1% polyethylene glycol (PEG) 400 (Thermo Fischer Scientific) in sterile PBS to achieve a concentration of 10 mM BIA. PPRX-1701 was prepared and generously provided by Cytodigm, Inc. as previously reported ^30^.

### G9-TCF reporter assay

G9-TCF cells were engineered by over-expressing a luciferase gene (Luc2) controlled by a TCF7-recognized promoter in the G9-PCDH cell line. Cells were plated in 96-well dark-well clear flat bottom plates at a density of 1500 cells/well. Next day, cells were treated with increasing BIA doses for 5 hours and then exposed to 10 µg/ml of D-Luciferin (Goldbio). Luminescence signal was quantified in the IVIS system.

### BIA quantification in vivo

G30-LRP cells were implanted in nude mice as previousy indicated, left to grow for 14 days, and injected with 20 mg/kg of BIA or PPRX-1701 (intraperitoneal). After 1 hour in circulation, mice were euthanized, perfused and tumor and brain tissue were harvested. Tissue was frozen at −80 degrees Celsius until processing. Quantification of BIA was performed using a Q-Exactive HFX Orbitrap mass spectrometer (LC-HRMS) (Thermo Fisher Scientific). Sample processing and analysis was performed as previously described ^30^.

### In vivo studies

For intra-cranial tumor implantation, GBM neuro-spheres were grown to 70% confluency before dissociated into single cell on the day of surgery. 50,000 cells were resuspended in 3 µl of sterile PBS and injected intracranially into the striatum (2 mm right hemisphere, 1 mm frontal, 3 mm depth from bregma) of mice under anesthesia and stereotactically fixed. Tumors were left to grow for approximately 2-3 weeks, depending the cell line. Animals were randomized to treatment groups. BIA injections consisted of 20 mg/kg (i.p.), except if indicated otherwise. PPRX-1701 was administered at 20 mg/kg (i.v.) via lateral tail-vein. Cisplatin injections were performed at 5 mg/kg (maximum tolerated dose, i.p.). All GBM tumor murine studies involved continuous condition and weight assessments, with endpoint considered when 20% of weight loss and/or moderate-to-high grimace scale and neurological symptoms were observed.

### ICP-MS for platinum quantification

Mice treated with cisplatin after BIA administration were euthanized, intra-cardially perfused with PBS and tissue harvested to be stored at −80 degrees Celsius. Tissue was processed and platinum (Pt195) was quantified using an Agilent 7900 ICP-MS, as previously described ^12^.

### Sodium fluorescein BTB permeability studies

Sodium fluorescein was administered to G30 tumor-bearing mice intravenously (i.v.) via lateral tail vein at 20 mg/kg. Then, 30 minutes after administration when peak fluorescence is reached in the brain, mice were euthanized for immediate brain tissue harvest. Fresh brain samples were visualized in Xenogen *in vivo* imaging system (IVIS). Quantification of pixel intensities from acquired images was performed in ImageJ. Tissue samples were then homogenized in 1 mL of 60% trichloroacetic acid in PBS and measured at 488 nm in a Molecular Devices SpectraMax M2 plate reader.

### Data and statistical analysis

Numerical results were analyzed, graphed, and statistically analyzed using the Prism software (GraphPad). Experiments were independently replicated at least three-times, unless indicated differently in the figure legends. Diagrams and workflow figures were generated using the BioRender software.

## Acknowledgements

This work was supported by NCI R01CA237063 (SEL) and NCI R21CA259734 (SEL 2023-2025 $250,000) grants. Choi-Fong Cho is supported by the R01 CA272573-01 (C-FC). Support for the ICP-MS instrumentation and analysis was provided by a core center grant P30-ES002109 from the National Institute of Environmental Health Sciences, National Institutes of Health. Bogdan Fedeles was supported by National Institutes of Health grants R01-CA080024, and P30-ES002109. The Thermo LC Orbitrap MS was supported by a National Science Foundation MRI award (1919870, K.D.P.) and the NIEHS Training in Environmental Pathology T32 program T32ES007272 (K.E.M.).

## Author contributions

J.L.J-M., and S.E.L. contributed to writing, data analysis and experimental conceptualization of this manuscript. Ay.Bh. and A.S.B. performed spatial transcriptomics data analysis from Ravi *et al*. (2022) manuscript. J.L.J-M., P.V-B., Z.X., W.H., N.M., J.R., M.O.N. and M.Z. performed animal experiments. J.L.J-M., P.V-B., M.F., A.S., Z.X., J.H., and M.S. performed western blots and immunofluorescence imaging from *in vitro* and *in vivo* samples. K.H. performed Luminex assay experiments and data acquisition. J.L.J-M and J.C. performed cell viability assays. C-F.C. and P.V-B. provided BBB spheroid usage and their data analysis expertise. K.E.M. and K.D.P. performed LC-MS and its analysis. B.I.F. provided training and analysis guidance for ICP-MS. B.W., W.L. and T.L. manufactured and kindly provided PPRX-1701.

