## Supplementary Data for "Modulation of blood-tumor barrier transcriptional programs improves intra-tumoral drug delivery and potentiates chemotherapy in GBM"

**A**

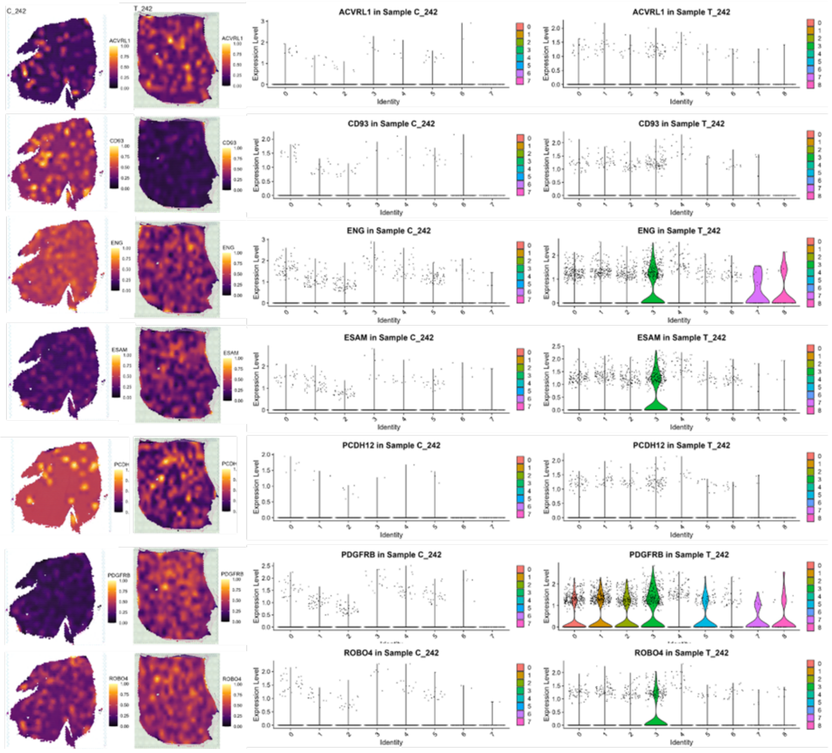

C_242
T_242
ACVRL1 in Sample C_242
ACVRL1 in Sample T_242
CD93 in Sample C_242
CD93 in Sample T_242
ENG in Sample C_242
ENG in Sample T_242
ESAM in Sample C_242
ESAM in Sample T_242
PCDH12 in Sample C_242
PCDH12 in Sample T_242
PDGFRB in Sample C_242
PDGFRB in Sample T_242
ROBO4 in Sample C_242
ROBO4 in Sample T_242
Expression Level
Identity


**B**

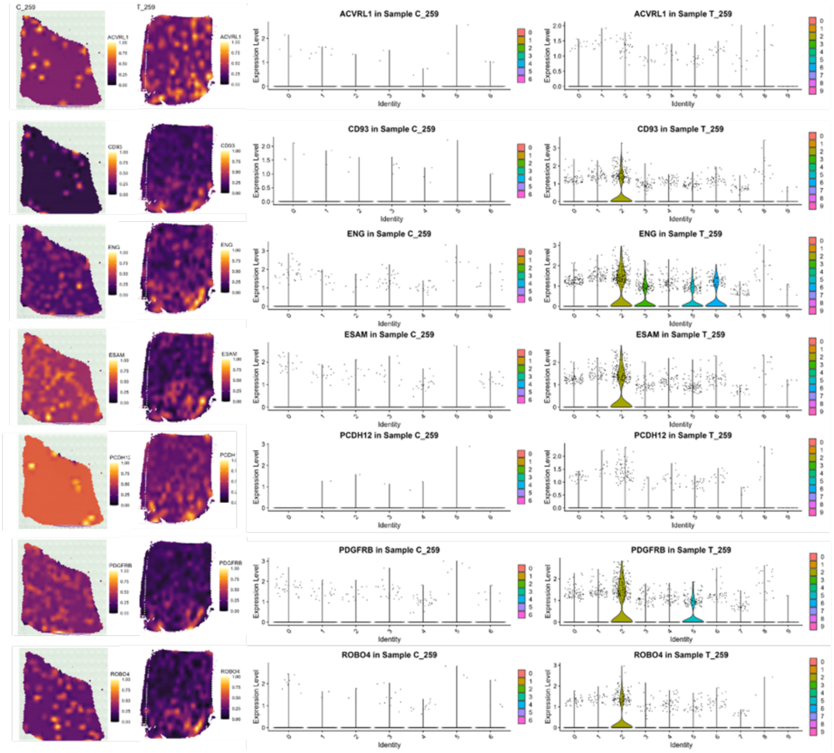

C_259
T_259
ACVRL1 in Sample C_259
ACVRL1 in Sample T_259
CD93 in Sample C_259
CD93 in Sample T_259
ENG in Sample C_259
ENG in Sample T_259
ESAM in Sample C_259
ESAM in Sample T_259
PCDH12 in Sample C_259
PCDH12 in Sample T_259
PDGFRB in Sample C_259
PDGFRB in Sample T_259
ROBO4 in Sample C_259
ROBO4 in Sample T_259
Expression Level
Identity

**C**

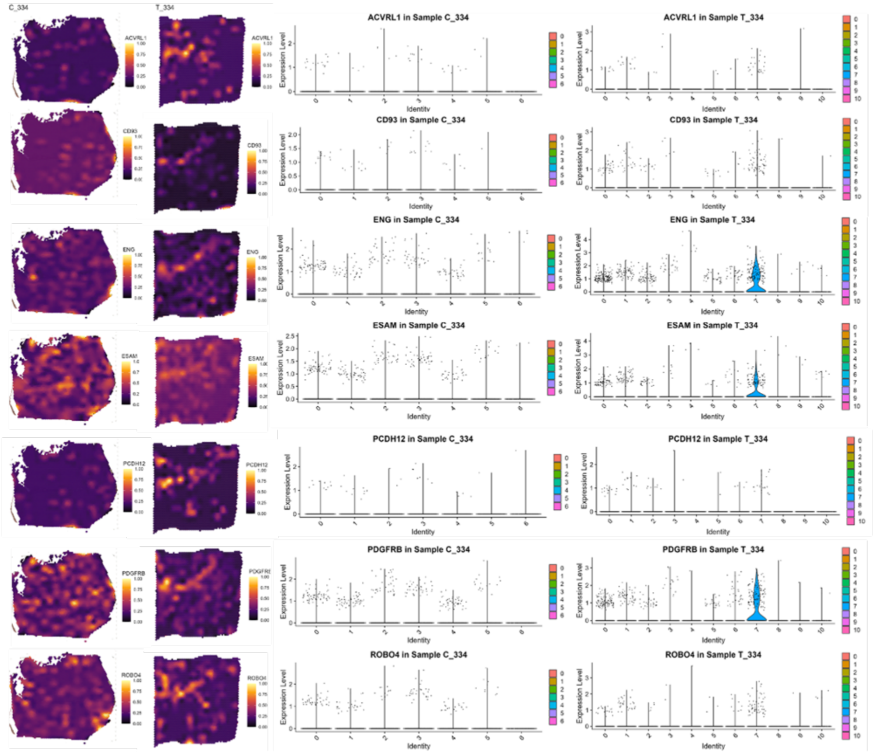


**Fig. S1. Spatial distribution of BTB-associated genes expression in GBM tissue compared to cortex control.** Surface plots (right) and clustered gene-expression violin plots (right) for cortex and tumor tissue from samples **(A)** UKF_242, **(B)** UKF_259, and **(C)** UKF_334.

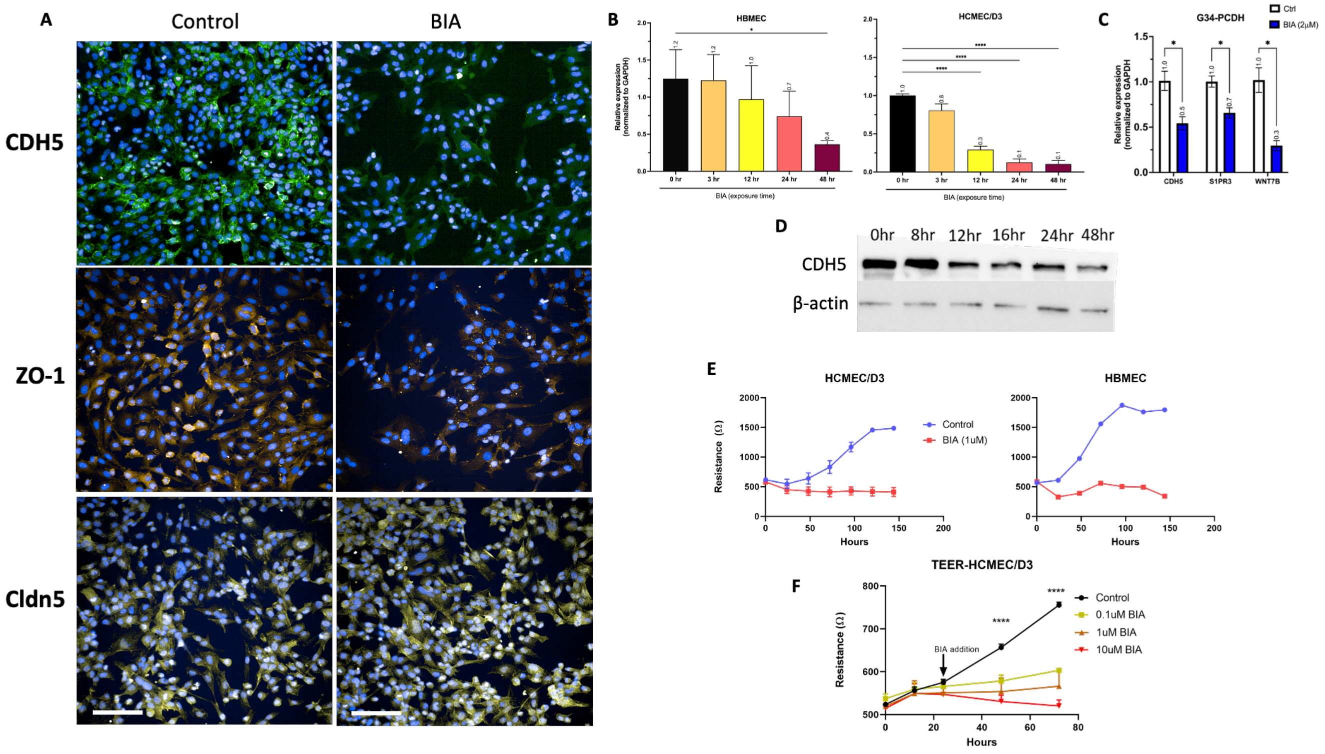


**Fig S2. BIA targets CDH5 and disrupts barrier formation of brain endothelial cells. (A)** IF staining of CDH5, ZO-1 and Claudin-5 in HCMEC/D3 cells treated with BIA (1 µM, 48 hours). Representative image is shown. Scale bar=100 µm. (**B**) Real-time PCR screening for CDH5 in HBMEC and HCMEC/D3 cells treated with 1 µM BIA for the indicated time-points. Mean and standard deviation are shown. Ordinary One-way ANOVA test * p=0.016, **** p<0.0001. (**C**) Real-time PCR screening for CDH5, S1PR3 and WNT7B in G34-PCDH neurosphere GBM cells treated with BIA (2 µM ) for 24 hours. Mean and standard deviation are shown. Multiple unpaired t-test was performed, * p<0.05. (**D**) Western blot analysis of CDH5 expression at different timepoints in HCMEC/D3 cells exposed to 1 µM BIA. Actin was used as loading control. (**E**) TEER analysis of HCMEC/D3 and HBMEC cells treated with BIA 24 hours after plating. Mean and standard deviation values are provided. (**F**) HCMEC/D3 dose response to different BIA doses analyzed by TEER. Two-way ANOVA test for control vs. BIA treatment comparisons (n=2/group), **** p<0.0001.

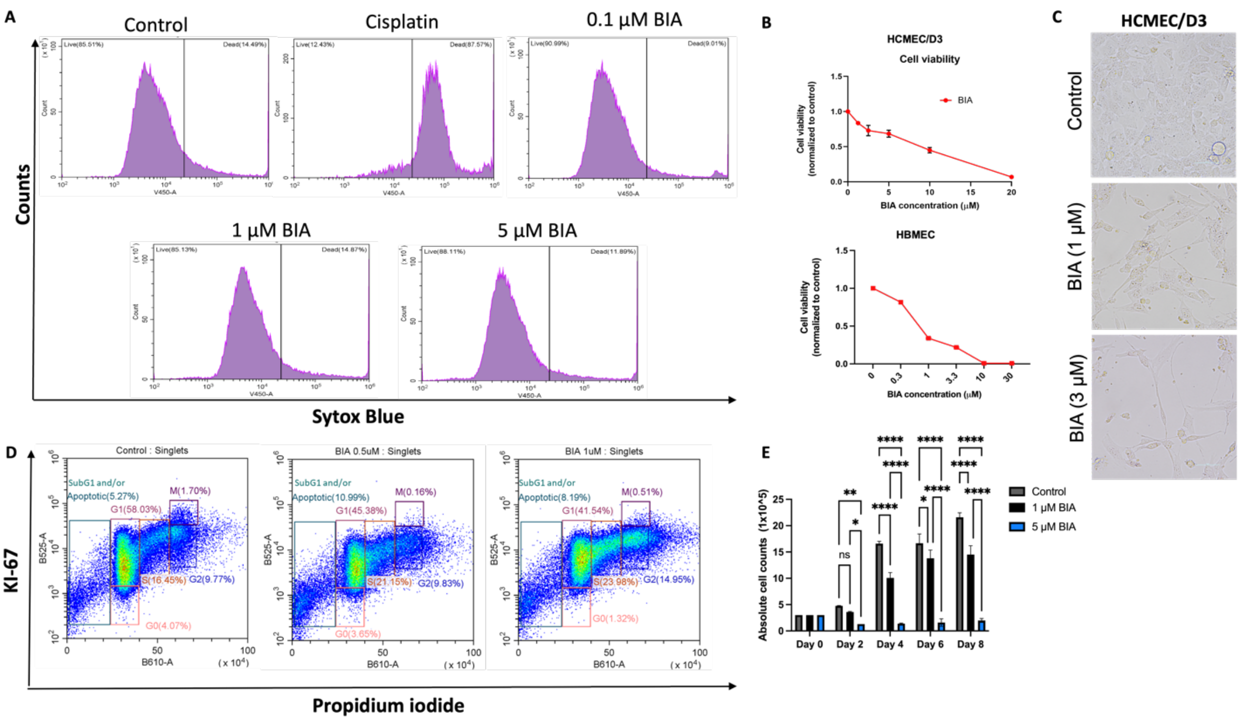


**Fig S3. BIA diminishes alters proliferation of brain endothelial cell lines but does not cause cellular death. (A)** Flow cytometry analysis of late apoptosis by Sytox Blue staining in HCMEC/D3 treated with different doses of BIA or Cisplatin (5 µM) for 72 hours. (**B**) Cell titer Glo viability assay on HCMEC/D3 and HBMEC cells treated with increasing concentrations of BIA for 72 hours. Mean and standard deviation of relative viability to control are shown, n=3 per group. (**C**) Brightfield images of HCMEC/D3 cells treated with BIA for 72 hours. Scale bar=50 µm. (**D**) Cell cycle analysis via flow cytometry of HCMEC/D3 cells treated with BIA (0.5 µM or 1 µM) or DMSO control for 72 hours. Representative figure of two-independent experiments is shown. (E) Absolute cell count numbers from HCMEC/D3 cells treated with BIA for the indicated durations and concentrations. Mean and standard deviation of duplicate independent counts are shown. Two-way ANOVA Tukey's multiple comparison test was performed. * $p<0.05$, ** $p=0.0030$, **** $p<0.0001$.

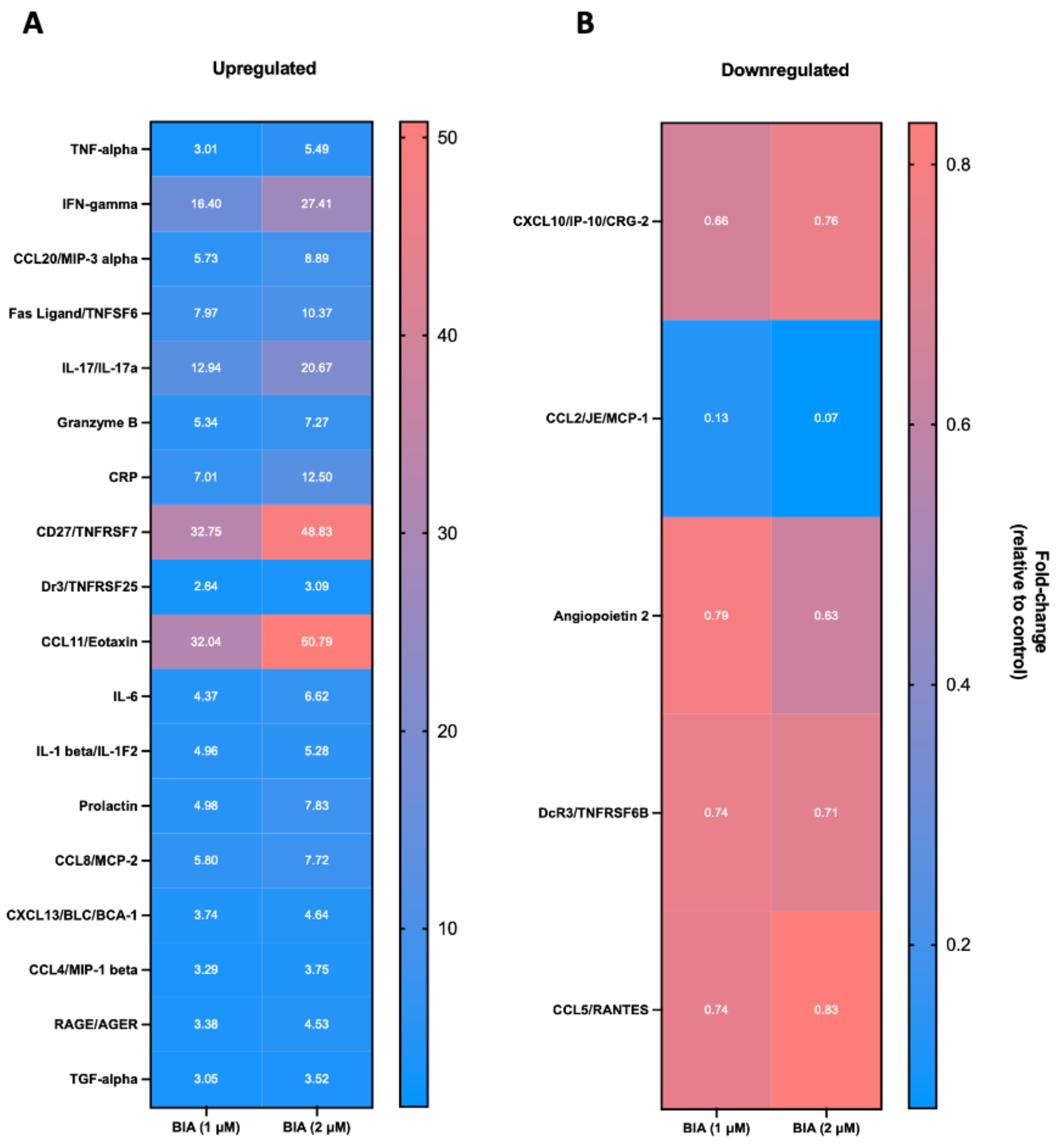


**Fig S4. Secretome changes after BIA exposure in brain endothelial cells.** Heatmap of Luminex cytokine analysis of HCMEC/D3 cells treated with 1 µM for 48 hours. (A) Fold-change relative to DMSO controls shows upregulated cytokines, and (**B**) shows fold-change decrease of cytokine secretion upon BIA doses.

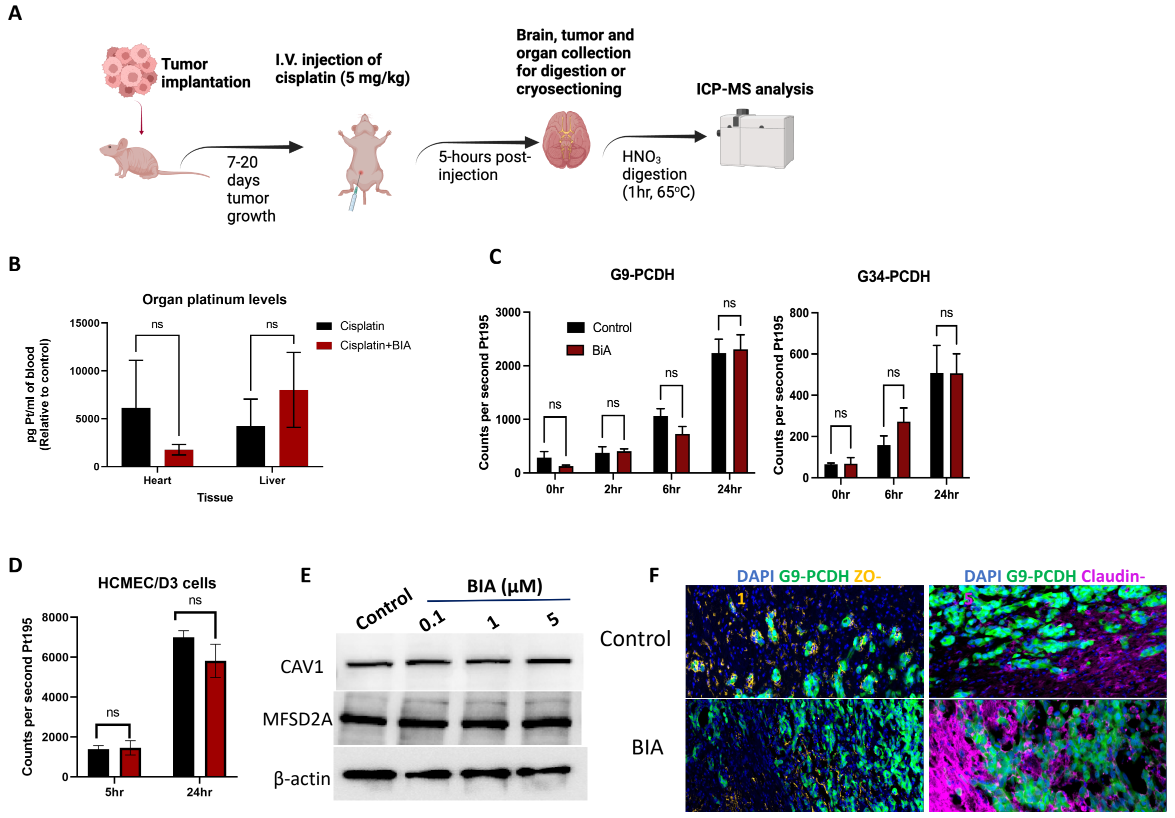


**Fig S5. BIA does not provoke direct intracellular cisplatin uptake in endothelial or tumor cells. (A)** Scheme of experimental layout for intra-cranial GBM tumor implantation in nude mice and subsequent treatment with BIA and cisplatin for ICP-MS quantification. (**B**) Platinum (Pt195) quantification in peripheral organs of cisplatin injected mice, 24 hours after BIA administration. Two-way ANOVA statistical analysis was performed from triplicate samples. (**C**) Platinum levels in G9-PCDH and G34-PCDH and (**D**) HCMEC/D3 cells after treated with 2 μM BIA and cisplatin for the indicated exposure times. Mean and standard deviation are shown from triplicates. Two-way ANOVA statistical analysis was performed. (**E**) Western blot from HCMEC/D3 cells treated with BIA at (1 μM ) for 48 hours. Actin was used as loading control. (**F**) IF staining of ZO-1 and Claudin-5 in the G9-PCDH xenograft model. Mice were treated with 20 mg/kg of BIA, and 24 hours later, brains were collected and fixed for sectioning and staining. Images taken at 20 x.

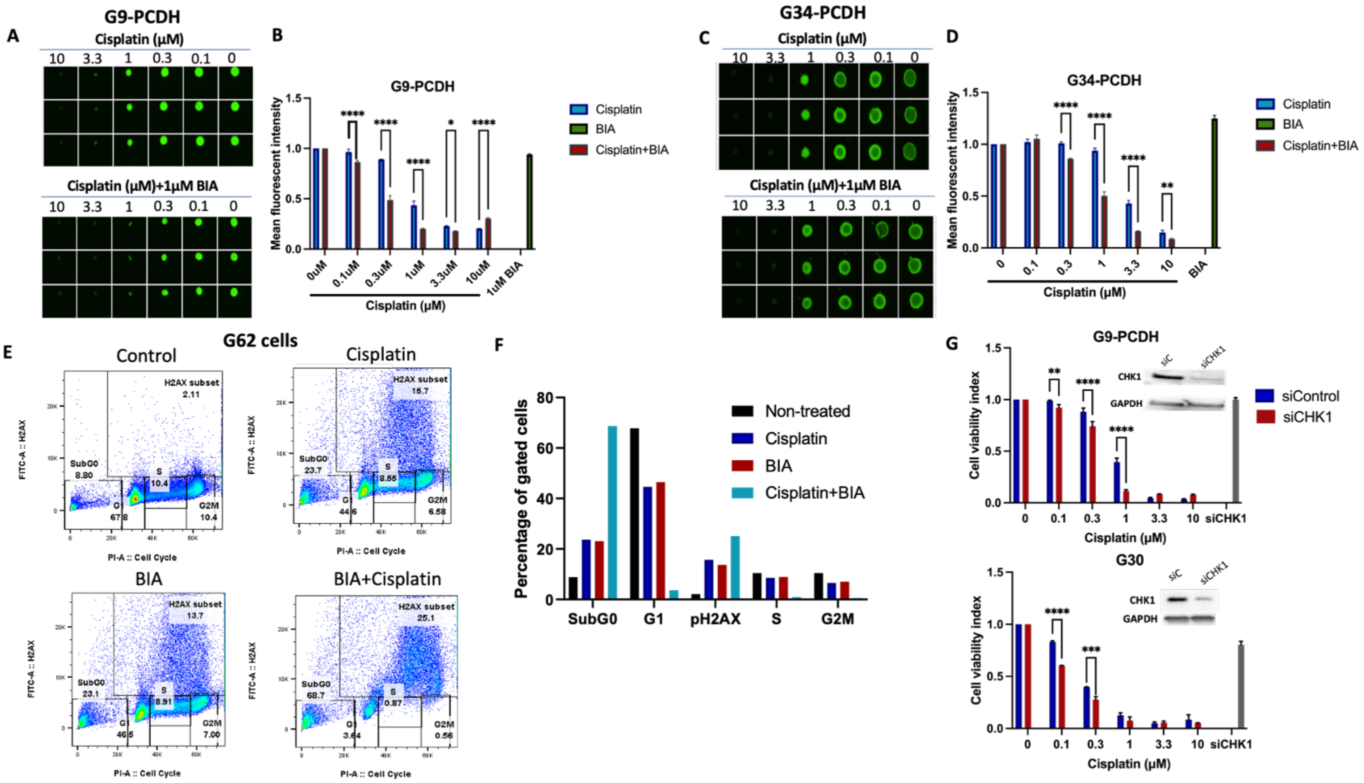

**Fig S6. Combinatorial treatment of BIA and cisplatin prevents neurosphere formation and increases DNA damage through CHK1 depletion.** Growth-in-low-attachment (GILA) assay for neurosphere formation assessment in (**A**) G9-PCDH and (**B**) G34-PCDH cell lines. Neurosphere size was measured using Image J (Fiji) in (**B**) and (**D**) for G9-PCDH and G34-PCDH, respectively. (**E**) Flow cytometry analysis of cell cycle by propidium iodide (PI) and DNA damage by γH2AX (Ser139) staining in G62 cells treated with BIA (1 µM) or cisplatin (1 µM) and their combination for 72 hours. Representative figure of two independent experiments is shown. (**F**) Graph of proportions of different cycle stages from data shown in (**E**). (**G**) Depletion of CHK1 via siRNA in G9-PCDH and G30 cells. Cells were then treated with different doses of cisplatin doses for 96 hours. Cell viability is shown relative to non-treated controls. Western blot confirmation of depletion is shown, with GAPDH as loading control. Mean and standard deviation are shown.

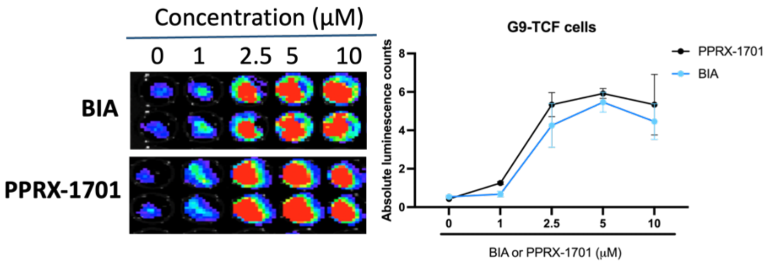


**Fig S7. PPRX-1701 BIA-loaded nanoparticles maintain GSK-3 inhibition potential and can reach brain and tumor tissues similar to BIA.** Luminescence detection of G9-TCF reporter cell line treated with increasing concentrations of BIA or PPRX-1701 nanoparticles for 5 hours. Luminescent signal was quantified by IVIS and shown in the graph to the right. No significant differences were found by Two-way ANOVA test, n=3.

| PECAM1 | CD34 | VWF | CLEC14A |
|---|---|---|---|
| ADGRL4 | DIPK2B | CLEC14A | TMEM204 |
| TES | ERG | FGD5 | GPR4 |
| ECSCR | MMRN2 | GPR4 | ESAM |
| CLEC1A | SCARF1 | ADGRF5 | DIPK2B |
| VASP | ACVRL1 | PDGFRB | ACVRL1 |
| GRAP | ARHGEF15 | FLT4 | VWF |
| FAM241A | FGD5 | DIPK2B | GJA4 |
| LXN | CD93 | PEAR1 | ADGRL4 |
| ARHGDIB | FLT4 | MMRN2 | DLL4 |
| CD40 | CDH5 | NPR1 | MMRN2 |
| TMEM255B | ADGRA2 | ADGRA2 | GRAP |
| GMFG | ADGRF5 | ARHGEF15 | ENPEP |
| ACVRL1 | PCDH12 | CDH5 | CDH5 |
| ICAM2 | CLEC14A | ROBO4 | ADGRF5 |
| CD93 | DLL4 | PCDH12 | RASIP1 |
| CAVIN3 | KDR | PDGFB | COX4I2 |
| ENPEP | BCL6 | ENPEP | FLT4 |
| ITGA1 | GIPC3 | ACVRL1 | CYYR1 |
| NOX4 | PEAR1 | TMEM204 | ARHGEF15 |
| DIPK2B | PDGFRB | ENG | PCDH12 |
| ESAM | RASIP1 | LAMC3 | LAMC3 |
| QPCT | ROBO4 | GIPC3 | ROBO4 |
| MYO1B | MYO1B | ERG | ECSCR |
| CLEC14A | UACA | SCARF1 | PDGFB |
| FAM162B | GPR4 | FAM43A | BCL6 |
| ENTPD1 | ENG | FLT1 | NR5A2 |
| GJA4 | LAMC3 | COL4A2 | HIGD1B |
| MYCT1 | MYCT1 | INSR | LRRC32 |
| TM4SF18 | VWF | CD93 | ERG |
| ERG | PRKCH | DLL4 | ITGA1 |
| MGST2 | ADCY4 | ITIH5 | MYCT1 |
| ATP8B1 | ENPEP | FOXC2 | SEMA3F |
| HIGD1B | GRAP | LRRC32 | RGS5 |
| MMRN1 | PECAM1 | HSPG2 | GIPC3 |
| STING1 | PDGFB | EXO3CL1 | ACE |
| KCNE3 | EOGT | KDR | EXOC3L1 |
| TNFRSF10D | ADGRL4 | FZD4 | FGD5 |
| PCDH12 | FLT1 | ADGRL4 | PEAR1 |
| LUM | ESAM | NID1 | NPR1 |
| CYYR1 | ECM1 | HRC | SOX17 |
| C1ORF54 | OR2A9P | RASIP1 | PECAM1 |
| ENG | EXOC3L1 | JAG2 | SCARF1 |
| FCMR | TIE1 | PAPSS2 | PDGFRB |
| KCNQ1 | HMCN1 | CD34 | NOTCH4 |
| CALHM5 | SOX17 | COL18A1 | PLXDC1 |
| FLVCR2 | HSPA12B | ITGA1 | CD34 |
| MPZL2 | TBXA2R | PRKCH | SH2D3C |
| LRRC32 | ACE | MYO1B | EGFL7 |
| MYL12A | NOS3 | BCL6 | NOX4 |
| FLI1 | CYYR1 | ZNF366 | ANGPT2 |

**Supplementary Table 1**. Co-expressed genes with PECAM1, CD34, VWF and CLEC14A in TCGA Firehose Legacy analyzed with cBIO.

| Co-expression with CDH5 |
|---|
| APOL1 |
| ATF4 |
| BMP2 |
| BMP7 |
| C3 |
| CCL2 |
| CCL5 |
| CCR7 |
| CDH1 |
| CDH2 |
| CDH3 |
| CDH4 |
| CDH7 |
| CEACAM1 |
| COL18A1 |
| COL1A1 |
| COL20A1 |
| COL5A3 |
| COL6A1 |
| COL6A2 |
| COL9A3 |
| DVL2 |
| E2F4 |
| ETHE1 |
| GFAP |
| ITGA2B |
| ITGA3 |
| ITGAE |
| ITGB2 |
| ITGB3 |
| ITGB4 |
| JUP |
| MAPT |
| MMP11 |
| MMP24 |
| MMP28 |
| MMP9 |
| NF1 |
| NR1D1 |
| OLIG2 |
| PCSK2 |
| PCSK4 |
| PECAM1 |
| PIEZO1 |
| PPARA |
| RBFOX3 |
| ROCK1 |
| RUNX1 |
| SLC6A4 |
| SMAD2 |
| SMAD4 |
| SMAD7 |
| SNAI1 |
| STAT3 |
| TFF3 |
| TGFB1 |
| TP53 |
| TUBB3 |
| WNT3 |
| WNT7B |
| WNT9B |

**Supplementary Table 2**. Genes co-expressed with CDH5 in GBM patient-samples from the spatial transcriptomics analysis in Ravi et al. (2022).

| Target | Forward primer sequence | Reverse primer sequence | Amplicon (bp) |
|---|---|---|---|
| *CDH5* | GGTCGATGCAGAGACAGGAG | GAGTCTCCAGGTTTCGCCA | 119 |
| *GAPDH* | CCAGCAAGAGCACAAGAGGA | ACATGGCAACTGTGAGGAGG | 104 |
| *S1PR3* | GACTGCTCTACCATCCTGCC | GATGCGTGCGTAGAGGATCA | 105 |
| *WNT7B* | GCGCTCGTCTCCGTCTATT | AGATGATGTTGGCTCCCAGG | 100 |

**Supplementary Table 3**. Table of primers used for real-time PCR analyses.
